# Dissecting structural and functional determinants of microtubule stabilization through guided chemical modulation

**DOI:** 10.1101/2025.10.23.684077

**Authors:** F Bonato, R París-Ogáyar, A Soliman, O Fernandez-Blanco, B Álvarez-Bernad, J Giménez-Abian, D Lucena-Agell, R Hortigüela, J Estévez-Gallego, W-S Fang, D Ondruskova, V Palomo, F Gago, M Braun, Z Lansky, JF Díaz, R Brandt, D Passarella, MA Oliva

**Affiliations:** Unit for the Development of Biological, Immunological and Chemical Drugs. Centro de Investigaciones Biológicas Margarita Salas – CSIC. Ramiro de Maeztu, 9. 28040, Madrid (Spain); Department of Chemistry, Università degli Studi di Milano, Via Golgi 19, 20133, Milan (Italy); Unit for the Development of Biological, Immunological and Chemical Drugs. IMDEA Nanociencia. Faraday, 9. 28049, Madrid (Spain); Department of Neurobiology, School of Biology/Chemistry, Osnabrück University, Osnabrück (Germany); State Key Laboratory of Bioactive Substances and Functions of Natural Medicines &MHC Key Laboratory of Biosynthesis of Natural Products, Institute of Materia Medica, Chinese Academy of Medical Sciences & Pekin Union Medical College, Beijing (China); Institute of Biotechnology, Czech Academy of Sciences, BIOCEV, Prague West (Czech Republic); Department of Biomedical Sciences and IQM-UAH Associate Unit. University of Alcalá, E-28805, Alcalá de Henares (Spain)

## Abstract

Paclitaxel (PTX) is a widely used chemotherapeutic, but its efficacy is limited by peripheral neuropathy, likely due to its structural impact on neuronal microtubules (MTs). To decouple MT stabilization from adverse structural changes, we designed, synthetized and characterized several PTX analogues. Our compound **1b** retains PTX’s stabilizing activity *in vitro* and in cells, and the **1b-**bound MTs preserve a native-like structure. These facts allowed us to investigate the influence of specific MT structural features on motor proteins behavior and interaction with MT-associated protein Tau. PTX-induced lattice expansion disrupted dynein-mediated retrograde transport and altered kinesin-1 motility. Additionally, PTX reduced Tau’s initial non-cooperative binding and envelopes growth rate while increasing dwell time and suppressing dynamic binding. In contrast, **1b** preserved more physiological Tau dynamics. These findings reveal that MT stabilization and structural modulation can be separated and highlight the functional importance of MT heterogeneity in maintaining neuronal transport and MAP interactions.

## INTRODUCTION

Microtubules (MTs) are dynamic cytoskeletal filaments essential for numerous cellular processes in eukaryotic cells. Formed by α/β-tubulin dimers arranged into protofilaments (PFs), they assemble into hollow cylinders with intrinsic polarity. MTs act as tracks for intracellular transport, play a central role in mitotic spindle assembly during cell division, and are main actors in neuronal development and cell motility. Their unique property, dynamic instability, is regulated by GTP hydrolysis, tubulin isotypes, post-translational modifications, and MT-associated proteins (MAPs), enabling rapid structural remodeling necessary for cellular functions. Because of their key role in mitosis, MTs are a major target in cancer chemotherapy. Drugs that interfere with MT dynamics disrupt cell division, leading to apoptosis in rapidly dividing cancer cells. Among these, paclitaxel (PTX), a taxane derived from *Taxus brevifolia*, is widely used to treat ovarian, breast, and lung cancers, and Kaposi’s sarcoma^1^. PTX stabilizes MTs by preventing depolymerization, thus disrupting cell division. However, its clinical use is often limited by dose-dependent neuropathy, a side effect likely due to MT functional alterations in neurons rather than off-target effects^2^.

MTs exhibit intrinsic structural flexibility. *In vitro*, tubulin assembles into filaments with 9–16 PFs, depending on the angle established among PF lateral interactions^3^. *In vivo*, the γ-tubulin ring complex enforces a conserved 13 PFs geometry, evolutionary favored for producing straight, stable MTs^4^. However, there are variations in PF number, such as those seen in platelets (14-15 PFs upon activation)^5^ or *C. elegans* mechanoreceptor neurons (15-PF MTs)^6^, where posttranslational modifications seem to play a crucial role^7^. Another critical structural feature is axial expansion, typically localized at MT tips and involved in MAP recognition^8^. Though this expansion is associated with stabilization due to the effect on MTs exposed to GTP and GDP analogs, taxanes or tubulin mutants^9–11^ it is also present in dynamic MTs^12^. Notably, PTX and related compounds can induce global lattice expansion^9,10,13^, potentially altering MAP interaction and contributing to neurotoxicity.

This study explores how taxane-induced MT structural alterations affect their function. Structural analysis of taxane-stabilized MTs using previously reported compounds revealed that certain C7-modified PTX derivatives can bypass the typical lattice expansion. By designing, synthesizing, and characterizing novel C7 derivatives, we engineered a compound that minimizes the MT structural distortion induced by MT stabilizing agents. Functional assays of intracellular transport and Tau interaction demonstrate that even subtle changes in MT architecture can significantly influence the interaction of these MAPs. These findings advance our understanding of how MT structural signaling modulates motors mediated transport and Tau binding, and they provide valuable insights for developing next-generation MT-stabilizing agents with enhanced efficacy and reduced side effects.

## RESULTS

### Paclitaxel derivatives at C7 override microtubule lattice expansion

We first examined whether MT lattice expansion is a general structural response to all taxanes^12^. To do this, we tested various taxane chemotypes using fluorescent PTX derivatives from our in-house library with modifications at the C2 position (2-debenzoyl-2-(*m*-aminobenzoyl)paclitaxel (2AP))^14^; C7 (Flutax2 (FTX2) and Hexaflutax (HFX))^15,16^, and C3’ (3’-N-m-aminobemzamido-3’-N-debenzamidopaclitaxel (3AP)^17^, (**Fig. 1A**). We analyzed the effects of these compounds on the structure of native MTs (assembled in the presence of GTP, thus GDP-bound with GTP caps) using X-ray fiber diffraction of aligned fibers^10^. This technique allows precise measurement of MTs structural parameters in solution. Briefly, the meridional profile (**Fig. S1, Fig. 1B**), provides information on the axial arrangement (i.e., compact vs. expanded lattice), while the equatorial profile (**Fig. 1C**) reflects lateral PF associations (i.e., MT diameter, PF number, and distribution of PF variants).

**Figure 1.**
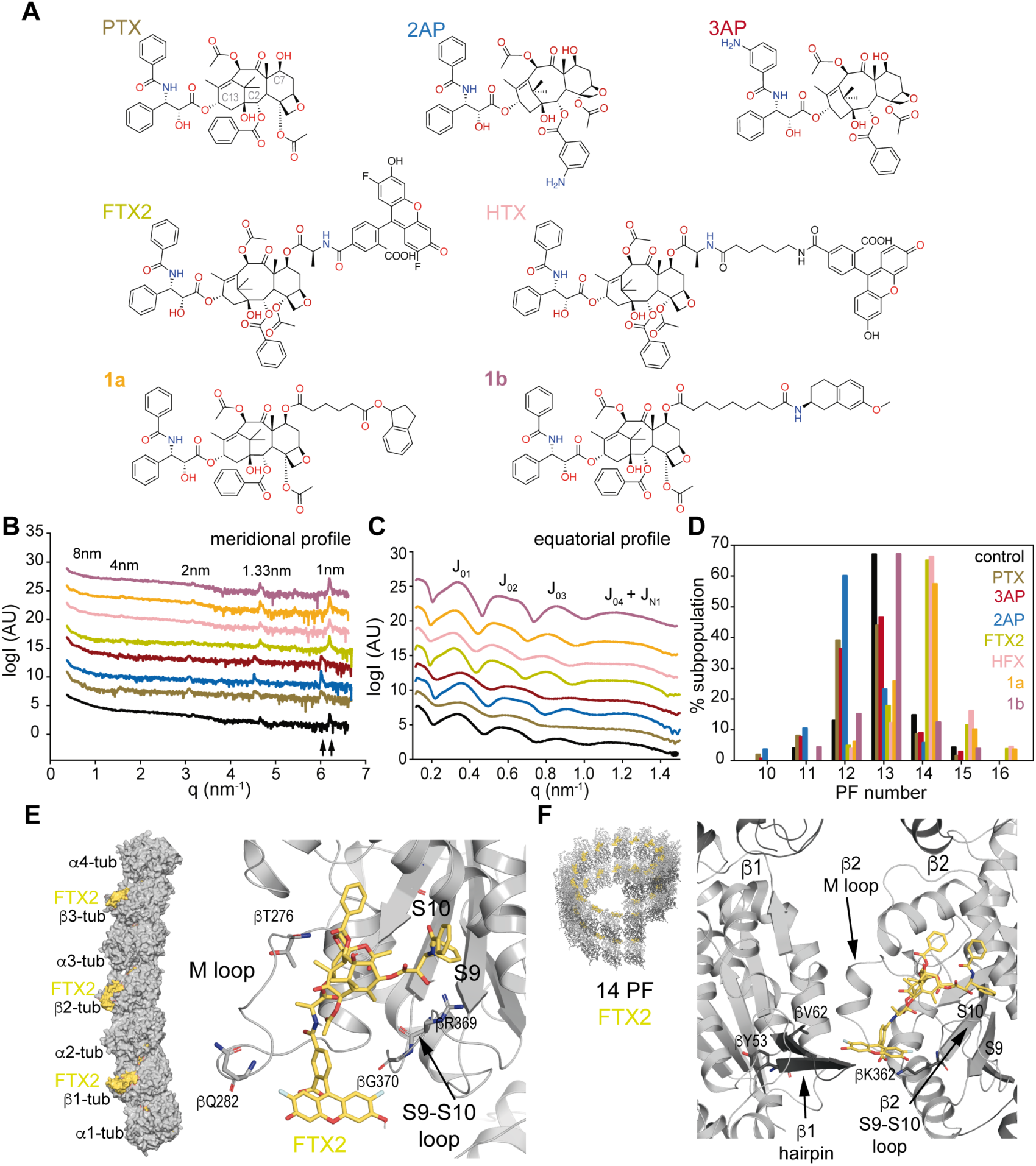
Molecules employed and structural analysis of related stabilized microtubules. **A.** Molecules used in this work. **B-C.** Fiber diffraction analysis; meridional **(B)** and equatorial **(C)** profiles of GTP-assembled MTs (GDP core and GTP cap), and GTP polymerized MTs stabilized with the different compounds. Profiles are shifted in the y axis for clarity and Bessel functions are labeled. In B, arrows point to the two possible positions of the 1 nm^-1^ layer band (compact vs. expanded lattices). **D**. Distribution of MT subpopulations according to the number of PFs in MTs assembled with GTP and MTs further stabilized with the compounds used in this study. **E.** Left. Surface representation of the PF used in MD simulations and built using three tubulin heterodimers highlighting FTX2 (yellow) in the taxane site of each β-tubulin. The PF is capped with an extra α-tubulin subunit to preserve β3-tubulin in a protein-protein axial contact context. Right. Zoom into the taxane site in β2-tubulin (cartoon representation) showing FTX2 (yellow, stick representation) orientation and highlighting main interacting residues from M and S9-S10 loops. **F.** Left. 14 PF MT used in MD simulations and Normal Modes and built as described in Methods section. Right. Zoom into the taxane site of one molecule (cartoon representation) showing FTX2 (yellow, stick representation) orientation and highlighting lateral interactions of fluorophore. Color code: control (black), PTX (brown), 2AP (blue), 3AP (red) FTX2 (yellow), HFX (salmon), **1a** (orange), **1b** (purple).

Consistent with prior studies, PTX induces axial lattice expansion of GDP-MTs^9,10^ as indicated by a peak position of the 1 nm^-1^ Bessel function at a *q* = 6.02 (monomer length of 4.17 nm in the real space, **Fig. 1B and Table 1**). Similarly, fluorescence-labeled PTX at C2 (2AP) and C3’ (3AP) induced lattice expansion. Surprisingly, both C7-derived compounds (FTX2 and HFX) promoted a compact lattice conformation (*q* = 6.20 nm^-1^, monomer length of 4.05 nm, **Fig. 1B**), thus overriding this otherwise consistent structural hallmark of taxane binding. Regarding lateral interactions, PTX and its C2 and C3’ derivatives led to the assembly of thinner MTs, whereas C7-modified compounds promoted wider filament formation (**Table 1**, **Fig. 1C**). In solution, native MTs primarily adopted a 13-PF configuration (> 65 %). PTX and 3AP induced a mixed population of 12-PF (∼ 40 %) and 13-PF (∼ 45 %) MTs, while 2AP skewed the population toward 12-PF MTs (∼ 60%). In contrast, C7 derivatives favored the formation of 14-PF MTs (∼ 65%, **Fig. 1D**). Neither FTX2 nor HTX could override the MT lattice expansion resulting from polymerization in the presence of non-hydrolysable GTP analog GMPCPP (**Table 1**, **Fig. S1 and S2**), suggesting that taxane and nucleotide binding sites modulate MT structure via independent mechanisms. Nevertheless, taxane-site stabilization in the presence of GMPCPP increased the average PF number (**Table 1, Fig. S2**) further supporting the proposed crosstalk between the nucleotide and the taxane sites^12^. Thus, PTX and 3AP moved toward a predominantly 13-PF population (∼ 65 %), while 2AP increased towards a more even distribution of 13-PF (∼ 40 %) and 14-PF (∼ 40 %) MTs. C7 derivatives shifted the population from 14-PF (55% FTX2 and 30 % HTX) to 15-PF (30 % FTX2 and 50 % HTX) MTs (**Fig. S2**).

**Table 1.** Fiber diffraction data analysis.

|  | Avg. radius<sup>1</sup> | Avg. PF number<sup>2</sup> | 1 nm<sup>-1</sup><br>band<sup>3</sup> | Avg. monomer rise |
| --- | --- | --- | --- | --- |
| GDP | 11.84±0.02 | 12.93±0.02 | 6.19±0.01 | 4.05±0.01 |
| PTX | 11.38±0.03 | 12.42±0.02 | 6.01±0.01 | 4.18±0.01 |
| 3AP | 11.42±0.03 | 12.46±0.03 | 6.01±0.01 | 4.18±0.01 |
| 2AP | 11.13±0.02 | 12.13±0.02 | 6.01± 0.01 | 4.18±0.01 |
| FTX2 | 12.57±0.02 | 13.74±0.02 | 6.19±0.01 | 4.06±0.01 |
| HTX | 12.77±0.03 | 13.96±0.03 | 6.20±0.01 | 4.05±0.01 |
| **1a** | 12.44±0.02 | 13.57±0.02 | 6.19±0.01 | 4.05±0.01 |
| **1b** | 11.75±0.01 | 12.82±0.01 | 6.19±0.01 | 4.06±0.01 |
| GMPCPP | 12.32±0.01 | 13.57±0.02 | 6.02±0.01 | 4.18±0.01 |
| PTX | 11.61±0.05 | 12.66±0.06 | 6.02±0.01 | 4.18±0.01 |
| 3AP | 11.94±0.02 | 13.03±0.03 | 6.02±0.01 | 4.17±0.01 |
| 2AP | 12.26±0.02 | 13.39±0.02 | 6.02±0.01 | 4.18±0.01 |
| FTX2 | 12.96±0.01 | 14.17±0.02 | 6.01±0.01 | 4.18±0.01 |
| HTX | 13.32±0.01 | 14.57±0.01 | 6.02±0.01 | 4.17±0.01 |
| **1a** | 13.07±0.02 | 14.29±0.02 | 6.02±0.01 | 4.17±0.01 |
| **1b** | 11.93±0.01 | 13.02±0.01 | 6.02±0.01 | 4.17±0.01 |

data are Mean±SEM
<sup>1</sup>nm as calculated from fitting J<sub>02</sub>
<sup>2</sup>calculated from interPF mean distance 5.7 nm<sup>56</sup>
<sup>3</sup>2pi nm<sup>-1</sup> as calculated from 1 nm layer line

### New C7-derivatives to tailor microtubule structural signaling

PTX structure-activity relationship studies have demonstrated that modifications at C7 are generally well tolerated, as the substitution of the 7-hydroxyl group does not significantly affect activity^18^. To investigate how the linker and bulky group at C7 influence lattice expansion and to explore chemical modulation of MT structure via the taxane site, we designed and synthetized two novel C7-modified compounds with tails of two lengths and distinct bulky groups: 2,3-dihydro-1H-indene (**1a**) and 6-methoxy-1,2,3,4-tetrahydronaphthalene (**1b**, **Fig. 1A** and Methods).

We first assessed their impact on the cellular microtubular network. Immunofluorescence staining in A549 lung carcinoma cells treated with either compound revealed hallmark features of PTX treatment, including MT bundling, loss of radial MT organization, and perinuclear clearing^19^ (**Fig. 2A**). In addition, cells exhibited aberrant nuclear morphologies, indicative of mitotic failure. These defects manifested as multinucleated cells resulting from multiple abnormal star-like spindles, leading to an aberrant arrangement of chromosomes. Notably, the concentrations required to induce these phenotypes were higher than for PTX (**Fig. 2A**). Cytotoxicity assays in A549 cells (**Table 2**) revealed that although both compounds retained submicromolar potency, PTX remained more potent. We then evaluated their ability to overcome intrinsic resistance mechanisms using cell lines overexpressing βIII-tubulin isotype (HeLaβIII)^20^ or P-glycoprotein, PgP, (Kb-V1)^21^, compared to parental HeLa and Kb-3.1 cells. Although these new compounds did not fully bypass these resistance mechanisms, they exhibited lower resistance-to-sensitivity (R/S) ratios than PTX, particularly in βIII-overexpression cells, which is consistent with previous *in vitro* experiments showing that PTX does not expand GDP-MTs assembled from recombinant α1β3 tubulin^22^. Interestingly, while compound **1b** was more potent in A549 cells, compound **1a,** exhibited a lower IC_50_ in HeLaβIII cells, suggesting that specific βIII-tubulin amino acid substitutions pattern may affect **1b** binding more strongly.

**Figure 2.**
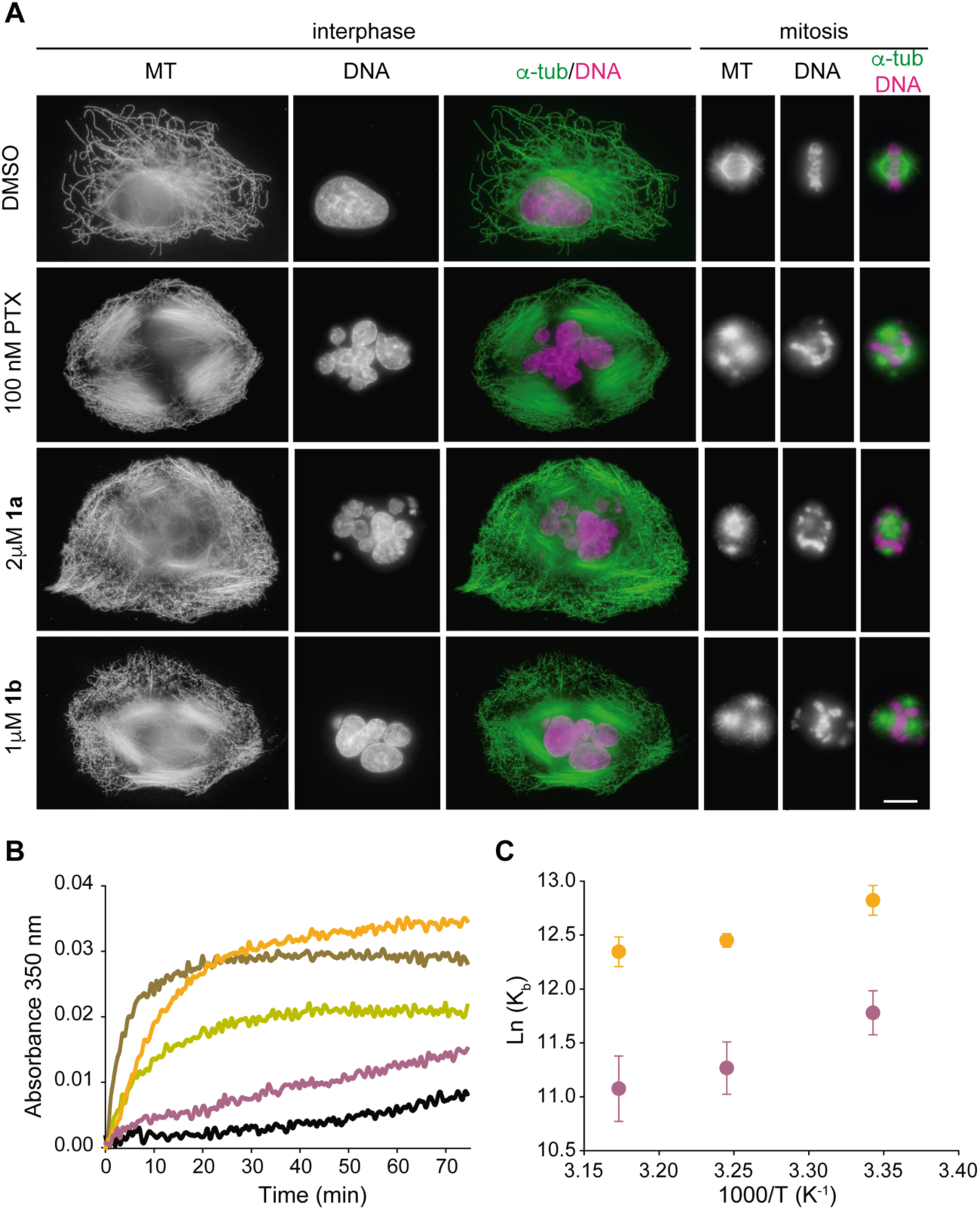
Cellular and biochemical characterization of compounds 1a and 1b. **A.** Effects of compounds on A549 cells. Each row shows α-tubulin immunostaining (DM1A antibody), DNA labeling (DAPI staining), and the merge of the previous ones (tubulin in green and DNA in magenta) of interphase cells (first three columns) and mitotic cells (following three). Top row panels are control cells treated with 0.5 % DMSO (vehicle), with the usual and evenly distributed microtubular network in interphase and normal bipolar spindle with chromosomes aligned at the metaphase plate during mitosis. In the following rows, 100 nM PTX, 2 μM **1a** and 1 μM **1b** treated cells, where the microtubular network is denser and with the presence of bundles during interphase, and a diversity of abnormal mitotic spindle containing misaligned chromosomes. All images are shown at the same magnification; scale bar 10 μm. **B.** Time course assembly of 25 μM tubulin in PEDTA buffer in the presence of the vehicle (DMSO, black line) or 27.5 μM of compound; PTX (brown), FTX2 (yellow), **1a** (orange), **1b** (purple). **C.** Van’t Hoff Plot of the equilibrium binding constants of compounds **1a** and **1b** at the assayed temperatures. Error bars represent the standard error of the measurement.

**Table 2.**
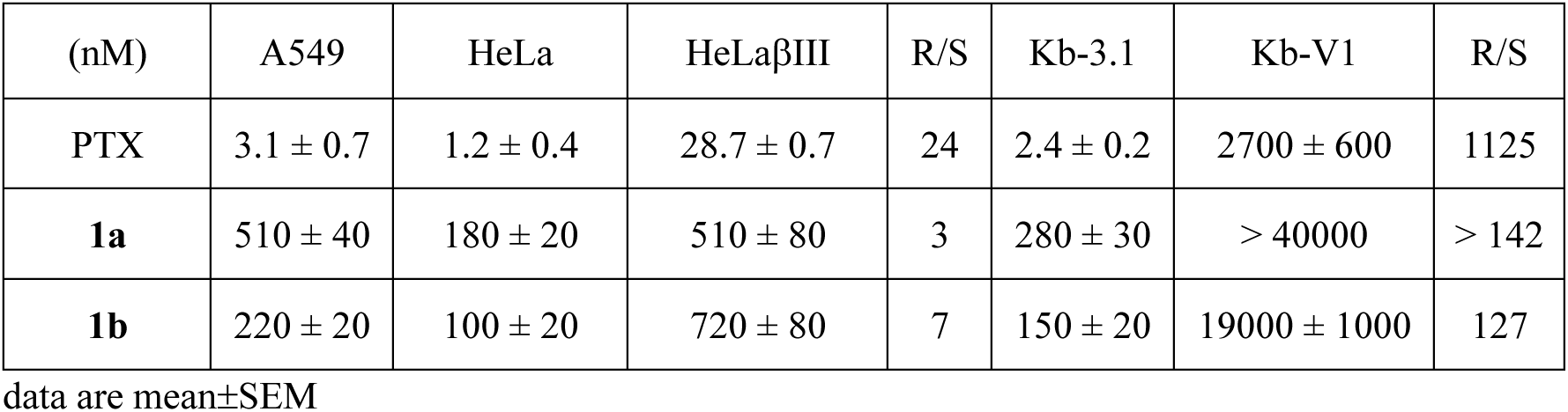
Comparative cytotoxicity (IC50) of new synthetized C7-derivatives.

Next, we evaluated the biochemical properties of these C7-modified compounds. *In vitro* polymerization assays revealed that the stabilization potency of **1a** was similar to that of PTX (**Fig. 2B**), whereas that of **1b** was lower and, similar to that of FTX2, which is probably indicative of higher cellular uptake for **1b** relative to **1a** (considering our cellular results, above). Consistently, **1a** exhibited ∼ 10-fold higher binding affinity for the taxane site than **1b** (**Table 3**), although the binding constants of both drugs was ∼ 100-fold lower than that of PTX^23^. Analysis of the thermodynamics of binding revealed that association reactions for both compounds were enthalpy-driven. Taxanes generally do not show significant changes in heat capacity upon binding^24^, and Vańt Hoff analysis (βH^0^_app_) confirmed that the binding of these new compounds was exothermic (**Fig. 2C**), consistent with PTX’s behavior (-51.4 ± 4.2 kJ mol^-1^)^23^. Overall, these results indicate that compounds **1a** and **1b** induce MT stabilization, and either the linker or the bulky moiety affects the interaction of the PTX core within the taxane site. Moreover, these results reinforce the premise that binding affinity does not directly predict a compound’s ability to promote MT assembly.

**Table 3.** Equilibrium binding constants and thermodynamic parameters of new synthetized C7-derivatives.

|  | **1a** | **1b** | PTX<sup>2</sup> | FTX2<sup>3</sup> |
| --- | --- | --- | --- | --- |
| K<sub>b</sub> (M<sup>-1</sup>) 26°C<sup>1</sup> | (3.7 ± 0.2) e5 | (1.3 ± 0.1) e5 | (2.64 ± 0.17) e7 | (5.9 ± 0.3) e7 |
| K<sub>b</sub> (M<sup>-1</sup>) 35°C<sup>1</sup> | (2.6 ± 0.1) e5 | (7.8 ± 0.8) e4 | (1.43 ± 0.17) e7 | (2.2 ± 0.2) e7 |
| $K_b$ ($M^{-1}$) 42°C<sup>1</sup> | $(2.3 \pm 0.1) \times 10^5$ | $(6.5 \pm 0.9) \times 10^4$ | $(0.94 \pm 0.23) \times 10^7$ | $(1.8 \pm 0.2) \times 10^7$ |
| $\Delta G^0$ app ($KJmol^{-1}$)<br>at 26°C | $-31.9 \pm 0.1$ | $-28.9 \pm 0.3$ | $-41.7 \pm 0.2$ | |
| $\Delta H^0$ app ($KJmol^{-1}$) | $-24 \pm 4$ | $-35 \pm 7$ | $-51.4 \pm 4.2$ | |
| $\Delta S^0$ app ($Jmol^{-1}K^{-1}$) | $-27 \pm 5$ | $20 \pm 5$ | $-29.3 \pm 13.1$ | |

<sup>1</sup>Data are the mean ± SEM values of three independent experiments with duplicates in each one
<sup>2</sup>as in<sup>79</sup>
<sup>3</sup>as in<sup>15</sup>. Equilibrium constants from anisotropy measurements.

To further dissect the contribution of the C7 substituent to binding, we calculated the incremental binding free energy (ΔΔG) associated with PTX-to-derivative modifications at 35 °C. The ΔΔG for **1a** (9.8 kJ mol^-1^) and **1b** (12.8 kJ mol^-1^) were similar to that of HTX (6.9 kJ mol^-1^)^25^, indicating that the C7 modifications introduced were relatively unfavorable. In contrast, FTX2 and in extent, also Flutax-1, which possess a ΔG^0^_app_ comparable to that of PTX, exhibited neutral to slightly favorable affinity change (-0.3 kJ mol^-1^ and-1.63 kJ mol^-1^)^15^. These findings denote that bulky and negatively charged groups confer more favorable interactions than smaller organic groups. The discrepancy between HTX and Flutax-1^15^, sharing the same fluorophore, may stem from differences in linker length (alanine aliphatic chain vs. single alanine). Combined with the observed differences between **1a** and **1b**, these findings suggest that shorter aliphatic chains yield higher binding affinities for the taxane site because the solvation of hydrophobic groups and longer aliphatic chains may need more energy.

Finally, we examined the structural effects of **1a** and **1b** on MT using X-ray fiber diffraction. Both compounds, consistently with other C7-derivatives, promoted a compact lattice structure, (**Fig. 1B**, **Table 1**). However, their effects on lateral PFs interactions differed. Compound **1a**, like FTX2 and HTX, promoted wider filaments with a dominant 14-PFs population (∼ 60 %, **Fig. 1D**). Remarkably, **1b** did not significantly alter the PF distribution relative to native MTs (> 65 % 13 PF). Nonetheless, neither compound was able to override the lattice expanding effect of GMPCPP (**Fig. S2**). Interestingly and differently from native conditions, compound **1b** did not show any alteration on the PF distribution in the presence of GMPCPP denoting that the interaction of this compound disable the crosstalk switch between the nucleotide and the taxane sites.

### Implication of C7 derivatives chains on protofilaments lateral interactions

To understand how lattice expansion is overridden by PTX C7 derivatives compounds we used computational methods. Initial molecular dynamics (MD) simulations of ligands FTX2 and **1b** in explicit water under comparable conditions revealed quite distinct conformational preferences (**Fig. S3A**) and a prevalence of poses incompatible with occupancy of the taxane site, in consonance with earlier work on other PTX-related taxanes^26^. Of note, FTX2 is shown to be able to form an intramolecular hydrogen bond involving the side-chain amide group; this interaction is not feasible in **1b** because the amide group has been separated further from the ester moiety attached to the taxane core by means of a polymethylene linker. This longer hydrophobic chain endows **1b** with much greater conformational freedom, which is translated into a more random orientation of the attached heteroaromatic ring relative to FTX2 (**Fig. S3A**) and correlates with its lower binding affinity.

We next analyzed the dynamic behavior of a fully solvated molecular model of a short PF consisting of three longitudinally αβ-tubulin dimers^12^, in both apo-and FTX2-bound forms (**Fig. 1E**). We found that, in the bound conformation, FTX2 maintains the intramolecular hydrogen bond reported above and places its fluorophore facing the intradimer α-tubulin subunit, sitting between the S9-S10 loop and the M loop (**Fig. 1E**). Consistent with our fiber diffraction data, simulations reliably reproduced a compact axial lattice within the dimer (α2:β2) and at the interdimer interface (β2:α3, **Fig. S3B**). The origin of lattice expansion has been attributed to the displacement of the S9-S10 loop, caused by backbone rearrangement at βR369-βG370, which propagates to the attached α-tubulin subunit^12^. Our MD trajectories show a shift in the O2’ taxane position compared to PTX and other taxanes structures. This change enables O2’ to act as a hydrogen bond acceptor from the amine backbone of βR369, rather than as a donor to the carbonyl backbone. Additionally, the oxetane group (O5) of the taxane hydrogen bonds to the backbone NH of βT276 (M loop), while the amide carbonyl (O4’) accepts a hydrogen bond from the (NE2)βH229 (central helix H7), in good agreement with previous crystal structures and MD trajectories^12^.

To understand the effect of ligand binding to lateral interactions between PF, we further studied MT patches (**Fig. 1F**). Binding of FTX2 to laterally assembled PFs invariably showed a strong ionic interaction between the free carboxylate of the benzoyl group attached to the fluorophore and the protonated amino group of βK362 (**Fig. 1F**). Thus fixed, the fluorinated xanthene moiety is positioned very close to the twisted β-hairpin (^53^YNEAAGNKYV^62^) of the β subunit in the vicinal PF. This means that FTX2 binding to MTs has an effect on lateral interactions that is more profound than that of PTX and analogues not bearing a long substituent at C7. In the case of **1b**, however, the longer and more flexible linker is capable of rotating quite freely both at the intradimer α1:β1 interface and at the interPF space (**Fig. S3C**).

Normal mode analyses of different 3-PF MT pieces herein described as [(α1:β1-α2:β2)/(α1’:β1’-α2’:β2’)/(α1”:β1”-α2”:β2”), **Fig S3D**] consistently revealed that the observed ligand-induced changes in PF number can be effectively approximated by the second non-trivial normal mode (**Movie S1**). Thus, depending on the direction (opening vs. closing motion), 13-PF MTs can evolve towards narrower 12-PF ones or towards wider MTs consisting of 14-15 PFs in a similar fashion as we showed before for laulimalide-and peloruside-bound MTs^3^. We posit that this low frequency motion can have a much larger amplitude than the spontaneous equilibrium thermal fluctuation when a ligand is bound because it is an activated process, *i.e*., ligand binding brings in additional energy that can stretch the structure along the normal mode much further and lock it in this state until ligand dissociation takes place.

### Different microtubule structural signals drive anterograde and retrograde intracellular trafficking

Building on our ability to chemically modulate MT structure, we next explored how these changes influence signaling patterns involved in MAPs interactions. Given that MTs are key players in intracellular transport – guiding organelles, vesicles and chromosomes via molecular motors like dyneins and kinesins – we assessed the impact of our compounds on transport dynamics in neuroblastoma SH-SY5Y cells. We combined live-cell imaging with previously developed tracking peptides (**Fig. 3A**): Cy5-DBP, which entails the sequence that specifically recognize Dynein Light Chain LC8 (retrograde transport), and Cy5-KBP, with a sequence that binds to the tetratricopeptide repeat of the kinesin-1 light chain KLC1 (anterograde transport)^27^. To ensure that our experimental conditions did not induce mitotic arrest or compromise cell viability, we applied high concentrations of the compounds (1 μM PTX or 2.5 μM FTX2, **1a** or **1b**) for a short incubation period (1 to 2.5 hours, including imaging time). We then assessed particle behavior by measuring mean track displacement and mean track velocity.

**Figure 3.**
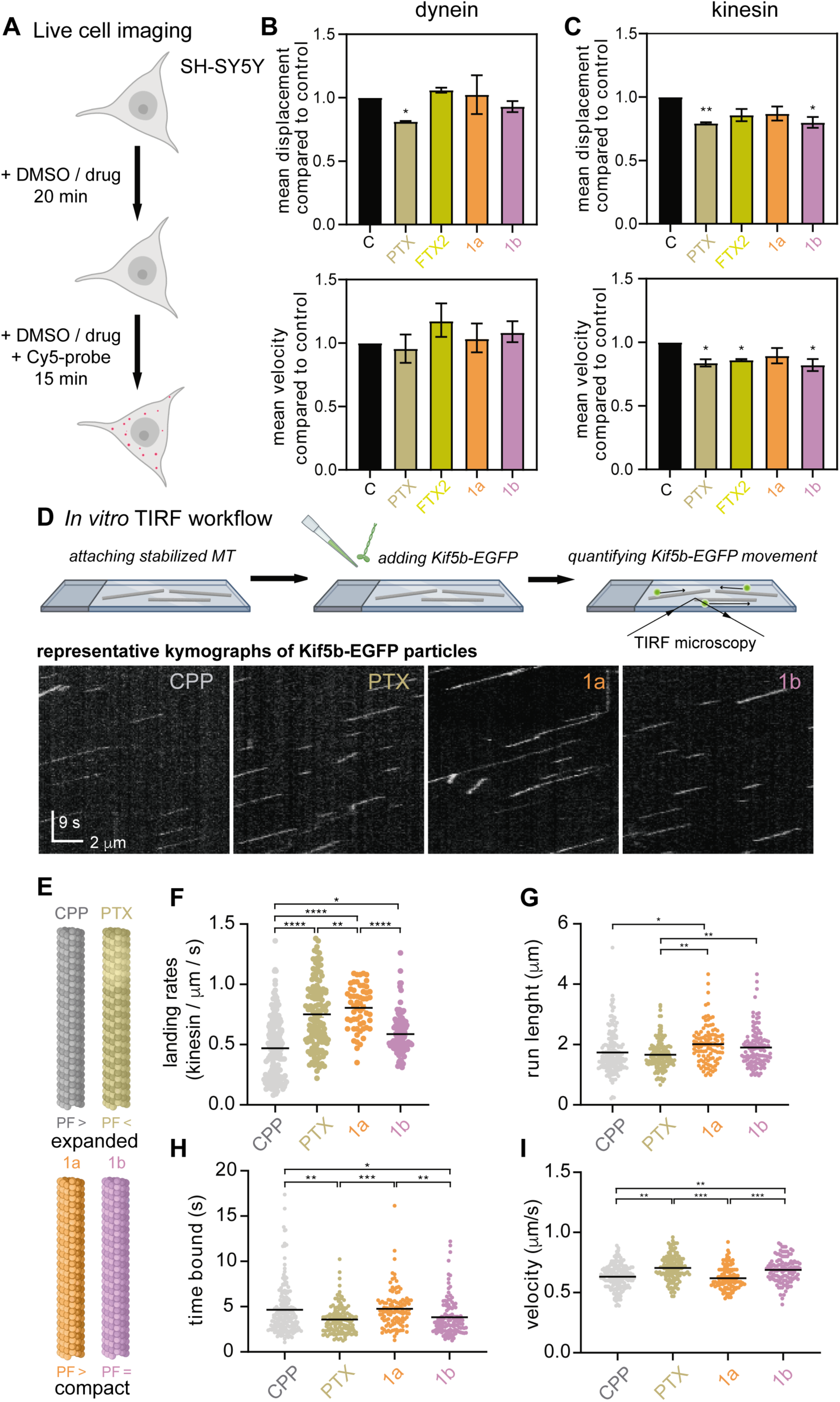
Characterization of motor proteins tracking. **A.** Schematic representation of live cell imaging of kinesin and dynein binding probes movement in SH-SY5Y cells. **B-C.** Particle movement parameters measured with Cy5-DBP (**B**) and Cy5-KBP (**C**) probes in the presence of 1 μM PTX or 2.5 μM FTX2, **1a** or **1b,** in non-differentiated SH-SY5Y cells. Data are the mean ± SEM values of two independent experiments with two to three different fields analyzed per experiment and statistical significance is as follows (** p < 0.01 and * p < 0.05). **D.** Schematic representation of the single-molecule kinesin-1 motility assay. Stabilized MTs were immobilized in the channel, Kif5b-EGFP were added and the motility of single motors was recorded using TIRF microscopy. Representative kymographs showing display the movement of multiple individual Kif5B-EGFP motor along a single MT; each line correspond to the trajectory of a single motor. Horizontal scale bar 2 μm; vertical scale bar 5 s. **E.** Simplified scheme of how studied compounds affect MT structure. **F-I.** Scatter plots and summary quantification of at least three independent experiments showing landing rate (kinesin/μm/s, **F**), run length (μm, **G**), time bound (s, **H**) and velocity (μm/s, **I**) of single Kif5b-EGFP molecules on MTs stabilized by different mechanisms. Each data point represents one motor molecule. Data pooled from n ≥ 2 experiments per condition across 12 MTs. Color code: PTX (brown), FTX2 (yellow), **1a** (orange), **1b** (purple), GMPCPP (gray)

Regarding the transport observed with Cy5-DBP, we found that only PTX had a significant effect, reducing particle displacement by approximately 20 % compared to control non-treated cells, without affecting speed (**Fig. 3B**, **Table S1**). These data suggest that dynein can sense lattice expansion, resulting in reduced contact time without compromising motor processivity. In contrast, we found no specific correlation between movement of Cy5-KBP and either axial or lateral MT lattice modifications (**Fig. 3C**, **Table S1**). Surprisingly, **1b** (which produces lattices close to the native state, **Fig. 1D**) led to reductions in kinesin binding probe tracking displacement (∼ 20 %) and velocity (∼ 16%, **Table S1**) similar to those observed with PTX (which causes lattice expansion and diameter reduction, **Fig. 1D**). Meanwhile, FTX2 (compact lattice with increased diameter) only reduced particle velocity slightly (∼ 14 %). However, **1a** (which induces similar lattice modifications to FTX2, **Fig. 1D**) did not affect particle tracking, which might be related to the lower cellular uptake observed (**Fig. 2, Tables 2**).

To gain mechanistic insight into the kinesin phenotype observed in cells, we reduced system complexity using an *in vitro* reconstitution assay. Specifically, we performed time-lapse imaging using total internal reflection fluorescence (TIRF) microscopy with immobilized and stabilized MTs in a flow chamber, into which we introduced 0.2 nM kinesin-1 motors (constitutively active human Kif5B 1-905-EGFP)^28^. We then analyzed individual kinesin movements to quantify landing rates (number of motors per unit MT length per unit time), run lengths, interaction times, and average velocities (**Fig. 3D, Movies S2-5**). In contrast to our experiments in cells, here we evaluated each structural modification individually to dissect the effects of MT lattice parameters on kinesin-1 mediated transport. Since non-stabilized MTs rapidly disassemble during the assay, we considered **1b**-stabilized MTs as the most suitable control because they exhibit a lattice structure close to the native form (**Table 1**). Each compound studied introduced distinct lattice alterations: **1a**-MTs increase the number of PFs in a compact lattice, and GMPCPP-MTs and PTX-MTs combine both axial expansion and an increase (GMPCPP) or decrease (PTX) in diameter (**Fig. 3E**).

In this simplified system, we observed that different kinesin motility patterns responded uniquely to the various lattice changes. Axial expansion (GMPCPP and PTX) did not correlate with time bound, velocity or landing rates (**Fig. 3H, I, F**). Larger MT diameter (GMPCPP and **1a**) increased the time bound (**Fig. 3H**) and decreased the velocity (**Fig. 3I**), while the landing rate did not correlate but decreased on GMPCPP compared with PTX (both expanded) while it increased on **1a** compared with **1b** (both compacted) (**Fig. 3F**). Lateral changes to the MT lattice did not significantly affect run length (**Fig. 3G**) as going slower (**Fig. 3I**) and staying longer (**Fig. 3H**) cancel each other out, and because the impact of axial expansion alone was minor, we conclude that individual lattice alterations are not primary determinants of kinesin motility. However, combined changes in axial spacing and PF number (e.g., **1b** vs. PTX; **1a** vs. PTX) lead to a more pronounced reduction in run length (**Fig. 3G**).

Contrary to previous reports, where the effect of lattice expansion was assessed by comparing GMPCPP stabilized MTs with non-stabilized GDP-MTs^29^, our results show that lattice expansion reduces kinesin-1 affinity for the MT lattice. This discrepancy may arise from the known structural heterogeneity of GDP-MTs (body and cap regions), whereas our system uses structurally homogeneous, stabilized lattices that enable us to isolate and evaluate the effect of discrete and combined structural features on kinesin-1 behavior. Our findings, further are at odds with assays demonstrating that gliding velocities of recombinant-single-tubulin-isotype MTs on kinesin-1 covered surfaces increase on expanded lattices^22^ and that kinesin-1 induces lattice expansion upon movement^30^. The discrepancy might result either from: (a) MTs assembled from mixtures of tubulin isotypes (as used in this work) responding differently to taxanes and GMPCPP than single-tubulin-isotype MTs; (b) synchronized teams of kinesin molecules behaving differently than single kinesins; or (c) from a combination of both. Notably, our findings further support the view that the expanded lattice states induced by GMPCPP and PTX – commonly used as MT-stabilizing agents *in vitro* – are structurally distinct^9,13,31^. This distinction should be carefully considered in future studies investigating MAPs-MT interactions.

### Both axial and lateral structural signals drive Tau binding and kinetics

Build on the established understanding that Tau’s dynamic interaction with axonal MTs is essential for modulating MT dynamics while preserving axonal transport^32,33^, we next investigated how MT lattice changes influence this interaction. To this end, we initially performed fluorescence decay after photoactivation (FDAP) experiments to assess potential changes in the interaction of the longest human CNS Tau isoform (Tau441) with MTs in axon-like processes of model neurons. We tagged Tau at the N-terminus with photoactivatable GFP (PAGFP) and expressed it exogenously in PC12 cells differentiated into a neuronal phenotype (**Fig. 4A**). After locally activating a subset of labeled Tau molecules in the middle of a process, we monitored FDAP over time in the activated region (**Fig. 4B**). Tau gradually dissipated from the activation site. Notably, only treatment with PTX led to a slower decay of the FDAP curves compared to control, indicating enhanced Tau binding to MTs in axon-like processes.

**Figure 4.**
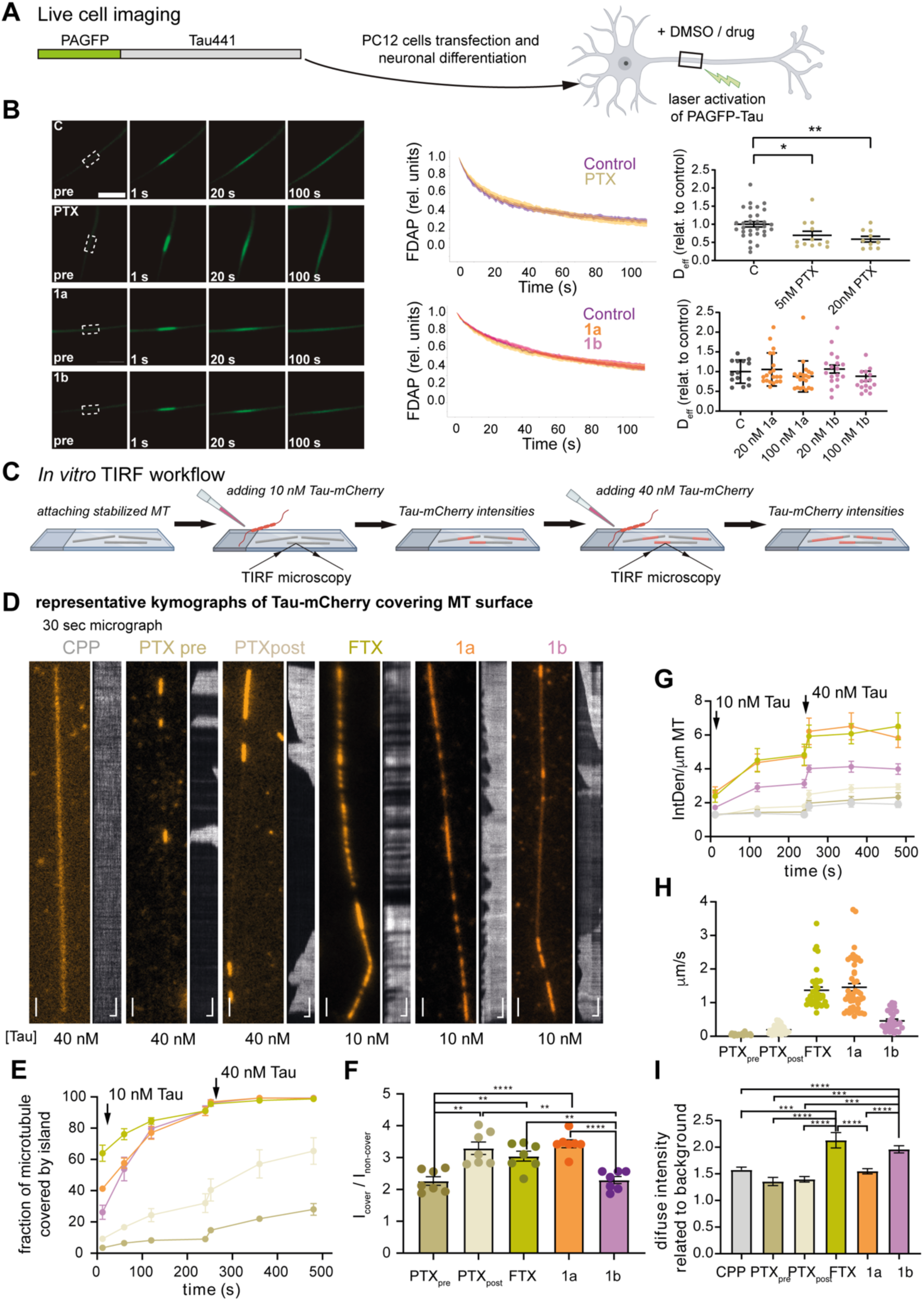
Characterization of Tau interaction with stabilized MTs. **A.** Live cell imaging of tau dynamics in axon-like processes of neuronally differentiated PC12 cells. Schematic representations of the expressed Tau construct with the N-terminal PAGFP fusion in green. **B.** Left: Representative time-lapse micrographs of a fluorescence decay after photoactivation (FDAP) experiment. Scale bar, 10 μm. The activation and recording region are indicated in the pre-activation image (6 μm length). Middle: FDAP diagrams after photoactivation of PAGFP-Tau-expressing PC12 cells. Mean ± SEM of Tau cells in the presence of PTX, compound **1a** and **1b** are shown. Right: Scatterplots of effective diffusion constants (D_eff_). Statistically significant differences between treatments as determined by unpaired Student’s t-tests with Welch correction are indicated. ** p < 0.01 and * p < 0.05. **C.** Schematic representation of TIRF microscopy assay setup. Label-free stabilized MTs are attached to a glass coverslip. 10 nM purified Tau-mCherry is added and its MT binding recorded in TIRF microscopy during 4 minutes. 40 nM Tau-mCherry is added and its MT binding recorded in TIRF microscopy for another 4 minutes. The fluorescence intensity of Tau-mCherry on each MT in the chamber is measured and analysed. **D.** Representative images (at frame 30 sec) and their corresponding kymographs of Tau-mCherry binding to MT stabilized with GMPCPP and PTX (40 nM Tau-mCherry) and, FTX2, **1a** and **1b** (10 nM Tau-mCherry). Vertical scale bars 2 μm and horizontal scale bars 60 s. **E.** Fraction of MT length covered by Tau envelopes after the addition of Tau-mCherry **F.** Mean ratio of intensity of covered regions divided by intensity of non-covered areas upon addition of Tau-mCherry on MTs stabilized with different drugs. **G.** Integrated density per MT μm after the addition of Tau-mCherry. **H.** velocity (μm/s) of envelope grow upon addition of 10 nM Tau-mCherry on MTs stabilized with different drugs. **I.** diffuse intensity signal related to background measured on MTs stabilized with different drugs. Error bars represent Standard Error of the Mean (SEM), n ≥ 2 experiments per condition, 12 MTs. Color code: PTX_pre_ (brown), PTX_post_ (light brown), FTX2 (yellow), **1a** (orange), **1b** (purple), GMPCPP (gray).

To quantify changes in Tau’s effective diffusion constant (D_eff_) as a result of its dynamic interactions with MTs, we measured FDAP at various PTX concentrations and fitted the resulting decay curves using a previously established reaction-diffusion model^34^. PTX treatment significantly reduced Tau’s D_eff_, with a notable effect already at 5 nM (**Fig. 4B**). This reduction corresponded to a ∼ 40% decrease in the unbound Tau population (4.4 % versus 7.5 % at 20 nM PTX), indicating increased MT binding. In contrast, neither compounds **1a** nor **1b**, even at concentrations up to 100 nM, significantly altered Tau-MT interaction (**Fig. 4B**). These findings suggest that an expanded lattice decreases Tau release dynamics in axon-like processes, while compounds **1a** and **1b** are neutral.

Previous studies *in vitro* have shown that Tau molecules strongly prefer compacted GDP lattices^35^. Tau binds cooperatively to MT surfaces, resulting in a cohesive layer depending on tau-tau interactions, termed MT envelope, which selectively regulates access to MT lattices, and shields MTs from severing enzymes^35–37^. On expanded PTX-MTs, Tau envelope also forms and the lattice can locally get compacted^35^. To further explore how MT lattice structure influences Tau binding, we employed an *in vitro* reconstitution assay. We deposited stabilized MTs in a flow chamber, and visualized them via interference reflection microscopy (**Fig. 4C**). We then added 10 nM Tau-mCherry (labelled purified Tau 441 isoform at the C-terminus with mCherry) and monitored its association with MTs using time-lapse TIRF microscopy. After 4 minutes, we increased Tau concentration to 40 nM, and continued imaging for an additional 4 minutes. We measured fluorescence intensities along MTs to quantify Tau binding (**Movies S6-17)**. Consistent with previous studies, Tau bound diffusively along GMPCPP-MTs^37^ and formed high-density envelopes that progressively expanded only on PTX-MTs^36,37^ (**Fig. 4D**). Even at the highest Tau concentration and observation time, we did not notice full MT coverage under these two conditions. Remarkably, we found differences between MTs assembled in the presence of PTX (PTX_pre_) and those stabilized with PTX after polymerization (PTX_post_). PTX_post_-MTs showed greater envelope coverage (up to 60 % at 40 nM Tau-mCherry compare to 30 % on PTX_pre_-MTs, **Fig. 4E**). PTX_post_-MTs also supported slightly faster envelope growth (0.20 ± 0.02 μm/s versus 0.054 ± 0.006 μm/s for PTX_pre_-MTs, **Fig. 4H**). Further fibre diffraction experiments under the same experimental conditions used for TIRF studies (i.e., MTs preparation 48 h beforehand) showed subtle structural differences among these stabilized MTs at the lateral interphase (**Table S2**). These findings suggest that MT lattice remodelling occurs slowly, at least in lateral PF arrangement, which likely contribute to the Tau binding patterns found.

We detected a markedly different Tau binding on compact lattices (i.e., FTX2-, **1a-**, and **1b**-MTs). At 40 nM, Tau-mCherry fully coated these stabilized MTs, further supporting that Tau interaction is favoured on compact lattices as previously described^35^. When we reduced the concentration to 10 nM, we could distinguish the two kinetic distinct phases of Tau binding: low-density regions with rapid turnover and high-density envelopes with slow turnover^36^ (**Fig. 4D-H**). At this lower concentration, PTX-MTs primarily displayed low-density Tau binding, with minimal high-density envelopes formation, especially on PTX_pre_-MTs (**Fig. 4E, G**). Therefore, to better capture Tau kinetics across all conditions, we used a two-step protocol: we first added 10 nM Tau-mCherry, followed by 40 nM. Under these conditions, Tau-mCherry bound similarly to FTX2-and **1a**-MTs, suggesting that Tau recognizes these alike lattices in the same way (**Fig. 4 D-G**). Interestingly, **1b**-MTs together with FTX2 showed higher diffuse fluorescence intensity than other stabilized MTs (**Fig. 4I**), likely indicating a more favourable non-cooperative and rapid turnover Tau binding. Otherwise, wider MTs promoted cooperative envelope formation, with an initial envelope coverage of 60 % (FTX2-MTs) and 40 % (**1a**-MTs) versus only 20% (**1b**-MTs, **Fig. 4E**)), and envelope grew rates of 1.37 ± 0.09 μm/s in FTX2-MTs and 1.5 ± 0.1 μm/s in **1a**-MTs compared to 0.46 ± 0.05 μm/s in **1b**-MTs (**Fig. 4H**). Since the cohesiveness of these envelopes depends on cooperative interactions through Tau N-^36^ and C-terminus^37^, the wider angle between PFs and also the different PF skews in FTX2-and **1a**-MTs might be favouring these interactions, though the specific mechanism remains elusive.

## DISCUSSION

PTX is one of the most relevant drugs in cancer therapy, but its use is hampered by the appearance of peripheral neuropathy, the most common dose-limiting side effect observed in patients. Although the precise mechanism underlying this pathology remains unclear, it is likely related to PTX’s interference with neuronal MTs^2^. Neuronal MTs are intrinsically more stable due to the expression of less dynamic tubulin isotypes and the presence of MAPs. Thus, PTX-induced stabilization may affect MT function differently in neurons. Main characteristic effects of the molecular mechanism of action of PTX is the structural distortion induced by axial lattice expansion and a reduction in PF number^9,10^. It is known that deviations from the canonical 13-PF geometry can induce PF supertwisting around the MTs^38^, and the conformational state of tubulin within MTs is known to modulate the interaction of MAPs such as VASH1/SVBP^39^ or doublecortin^40^ Given that the MT structure serves as a key signalling platform for cellular recruitment of motor proteins and MAPs, it becomes evident that such structural distortions, which in addition spread along the fibre, may underlie MT malfunction by altering their recognition patterns in axons following PTX exposure.

To test this hypothesis, we set out to design a compound capable of exerting PTX-like activity without inducing structural distortion of MTs, and to investigate the cellular effects of such compounds. Based on our initial X-ray fibre diffraction results – and considering that modifications at the C7 position of PTX have shown potential to modulate its biological activity^41^ – we designed, synthetized and characterized novel compounds that allowed us to specifically tailor MT structure. These compounds retained the ability to stabilize MTs both *in vitro* and in cells, although with reduced potency, which we then correlated with differences in linker length and attached moieties. Notably, compound **1b** emerged as the only derivative able to mimic a native-like MT structure while maintaining MT stabilization activity. This finding prompted us to explore which structural features of MT drive MAP recognition. Our *in vitro* characterization provided the best up-to-date mechanistic insight, while cellular studies integrated this knowledge into a more complex and heterogeneous context, shaped by drug-specific parameters (resulting in non-equimolar intracellular concentrations), the “tubulin code” – such as variations in tubulin isotypes that also regulate the PF number^42^ – and the presence of additional MAPs. Because stabilized MTs exhibit quasi-homogeneous structures, our approach enabled the identification of how individual structural cues affect intracellular transport and modulate the behavior of axon-specific MAPs like Tau.

Neurons critically depend on efficient intracellular transport, which is essential for cargo trafficking from the soma to synaptic terminals (via kinesin-mediated anterograde transport) and in the reverse direction (via dynein-mediated retrograde transport). Our data in SH-SY5Y neuroblastoma cells reveal a distinct effect of MT lattice expansion on retrograde transport, particularly in reducing the distance measured by the dynein binding probe, suggesting that PTX may impair the intracellular trafficking. In contrast, kinesin-1-binding probe appears less sensitive to lattice expansion and more influenced by other structural features. Unexpectedly, compound **1b** also reduced both length and speed of kinesin-1 binding probe transport, possibly due to enforced MT homogeneity in cells, where MTs typically display dynamic, asymmetric structural fluctuations^43^. The similarity of this effect to PTX treatment highlights the complex regulation of kinesin-1 movement by MT lattice features. Differences from our previous results in A549 cell^27^ may arise from differences in the tubulin code (i.e., tubulin isotypes expression and post-translational modifications), which are known to influence motor interactions with MTs^44^, as well as differential drug uptake and, drug’s binding affinity to tubulin and MTs. In addition, we have found significant differences with previously reported PTX-induced increases in retrograde mitochondria transport and enhanced anterograde velocity^2^. This study used lentiviral transduction of Mito-dsRed to label mitochondria, which in combination with differences in cell lines, PTX exposure duration, and dosage could also account for the observed discrepancies. Furthermore, the mobilization of mitochondria is mediated by multiple motors simultaneously. Recent *in vitro* assays demonstrated increased gliding velocities of PTX and GMPCPP-MTs (expanded-lattices) propelled by synchronized teams of kinesin-1 motors^22^. The mixed results provided by our kinesin-1 single molecule stepping assays (decreased velocity on PTX-MTs while increased on GMPCPP-MTs) indicate that synchronized teams of motors, for example propelling organelle movement *in vivo*, might be affected differently than single kinesin molecules. Follow-up *in vitro* experiments further clarified how distinct structural changes influence kinesin-1 binding and motility: lattice expansion decreases landing rates, while alteration on the PF number, which is a feature already described on axonal MTs^45^, enhances kinesin-1 affinity. However, when both axial and lateral modifications are present, the effects become more complex. Compared to native-like MTs, PTX-stabilized lattices reduce run length (in line with cellular observations, although velocity is unaffected *in vitro*).

Tau binding to MTs is generally considered not to strongly impact axonal transport rates^46^, likely due to its “kiss-and-hop” dynamics ^33^. Our results in differentiated PC12 neuronal cells demonstrate that PTX-induced lattice expansion enhances Tau binding to MTs, which in turn has the potential to impair axonal transport^32,33^. Using a simplified *in vitro* system, we uncovered the structural basis of Tau-MT recognition. We show that Tau envelope formation arises from specific lattice features that promote cooperative Tau assembly once nucleation occurs. Moreover, our data suggest that Tau senses both axial and lateral structural cues, which shape its binding kinetics and, in turn, influence transport regulation and other MAPs interaction^35–37^. Importantly, Tau preferentially recognizes compressed lattices. While envelope formation is still possible on expanded lattices, it growth rate is markedly slower, consistent with direct competition between PTX and Tau binding^47^. Taken together, our *in vitro* and cellular findings indicate that on PTX-stabilized MTs, Tau’s initial affinity decreases, but its dwell time once bound increases, likely due to PTX release^47^, resulting in reduced kiss-and-hop dynamics.

In conclusion, our study underscores the functional relevance of MT heterogeneity *in vivo*, a key determinant of the diverse biological processes mediated by the cytoskeleton. This diversity should be carefully considered in the context of pharmacological intervention. At the same time, we demonstrate that our compound **1b** preserves PTX’s stabilizing effects on tubulin and MT, though with lower potency, while maintaining native lattice architecture. These results establish that taxane-induced polymerization and lattice remodeling can be uncoupled, opening new avenues for rationally tuning MT architecture through chemical design.

## MATERIALS AND METHODS

### Protein and chemicals

Purified calf brain tubulin and chemicals were obtained as previously described^24^. Tau-mCherry was expressed and purified as described previously^28^. Human Kif5B-EGFP (amino acids 1–905), containing a C terminal fluorescent-tag followed by a 3C protease cleavage site and a 6 × His-tag was expressed and purified as in^48^ with some modifications. Prior to purification, cells were resuspended in 1:1 ratio of lysis buffer (30 mM HEPES pH 7.5, 300 mM NaCl, 0.5 mM ATP, 5% glycerol, 20 mM imidazole, 0.5 M TCEP) supplemented with 1x Protease inhibitor cocktail and benzonase to the final concentration of 25 units/ml. Cells were homogenized by repeated pipetting. Cells were lysed by spinning for 45 min at 40,000 rpm in 45 Ti fixed-angle rotor (Beckman Coulter) at 4 °C. The supernatant was incubated with pre-equilibrated HisPUR NiNTA agarose resin (Thermo Fisher Scientific) for 1 h in a tightly closed gravity-flow column (Thermo Fisher Scientific) at 4 °C by slowly rotating. Subsequently, the resin was washed once lysis buffer and then with buffer B (30 mM HEPES pH 7.5, 300 mM NaCl, 0.5 mM ATP, 5% glycerol, 30 mM imidazole, 0.5 M TCEP). The protein was eluted from the resin after an overnight cleavage of the 6 × His-tag with 3C HRV protease in lysis buffer at 4 °C. The eluted protein was further purified via size exclusion chromatography on Superdex® 200 10/300 GL column equilibrated in exclusion buffer (30 mM HEPES pH 7.5, 300 mM NaCl, 0.5 mM ATP, 5% glycerol, 0.5 M TCEP). Peak fractions were analyzed via SDS-PAGE, pooled and the protein was concentrated with Amicon Ultra-15 Centrifugal Filter tubes (Millipore). Purified proteins were aliquoted, snap-frozen in liquid N_2_, and stored at – 80 °C.

#### Fiber diffraction

X-ray fiber diffraction data were collected in beamline BL11-NDC-SWEET at ALBA Synchrotron. The energy of the incident photons was 15 KeV or equivalently a wavelength, l, of 0.827 Å^10^. Purified bovine brain tubulin was diluted to a final concentration of 100 μM in FD buffer (0.8 M Pipes-K, 1 mM EGTA, 0.2 mM Tris, 1 mM DTT, 3 mM MgCl_2_ pH 6.8) and incubated on ice for 10 min. Insoluble aggregates were removed by centrifugation at 13,500 rpm for 10 min at 4 °C using Spin-X® 0.45 μM cellulose acetate filters (Costar, Corning, USA). Tubulin was then supplemented with 2 mM GTP or 0.5 mM GMPCPP and 200 μM of the tested ligands. All samples were incubated for 30 min at 37 °C to induce the maximum fraction of polymerized tubulin and then, mixed in a 1:1 volume ratio with FD buffer containing 2% methylcellulose (MO512; Sigma-Aldrich). Samples were centrifuged 10 s at 2,000 g to eliminate air bubbles and transferred to the space between a mica disc and a ring-shaped cover slip arranged in the shear-flow device^49^. The device was kept at 37 °C during measurements and the quartz disc rotates at 10 r.p.s to achieve the shear flow alignment. The SAXS diffraction patterns were collected in a Dectris PILATUS3S-1M, with an active area of 168.7 x 179.4 mm^2^, an effective pixel size of 172 x 172 mm^2^ and a dynamic range of 20 bits. Background images were acquired in the same conditions, using PM buffer with methylcellulose without tubulin. Four diffraction images per sample were acquired, each with a 150 s exposure, as higher frame numbers risked radiation damage. For each condition, 8–12 images were collected across 2–3 independent experiments.

##### Data processing

The whole analyses were performed using custom routines implemented in Python, leveraging libraries such as *NumPy*^50^, *pandas*, and *SciPy*^51^. Raw fibre diffraction images were first corrected by subtracting an averaged background image. For each condition, multiple buffer images were normalized and averaged to generate a representative background signal that was subtracted pixel-wise from each sample image to isolate the scattering signal of the sample. This pre-processing step eliminates parasitic scattering contributions and enhances the contrast of the diffractograms. Subsequently, images were subjected to azimuthal integration using the pyFAI library^52^ and a previously calibrated geometry file (poni file), containing detector distance, beam centre, and tilt angles, and obtained via standard calibration images with silver behenate. Integration was carried out in specific azimuthal sectors centred around the equatorial and meridional axes. To improve signal quality along the equatorial axis, we also subtracted the residual background scattering from non-oriented species (mainly unassembled tubulin). For doing this, we considered two off-axis regions located at ± 45° relative to the equator because at these areas the scattering from the oriented species is expected to be negligible. Polarization correction and solid angle normalization were applied during the integration. The resulting profiles provided the scattering intensity as a function of the scattering vector q ([4π sin(θ)] / λ), with separate curves for equatorial and meridional directions. Finally, 1D scattering profiles were analyzed to determine main structural parameters of the fiber assemblies. Equatorial profiles were initially fitted using a diffraction model based on previous approaches^53–55^ of the form I(q) = wₙ · ([J₀(q·rₘ) · F(U)]² + fₙ · [Jₙ(q·rₘ) · F(U)]²) and hence, considering the zero^th^-and n^th^-order Bessel functions of the first kind, which represent the two helical scattering components of the MT lattice. However, we found that using J_0_ and J_n_ the fitting was not satisfactory because only applied to minimal regions of the profile. Considering that J_01_ is affected by central scattering and J_03_ could be convoluted with J_n_, we selected J_02_ for approaching a new fitting. Because determination of the PF distribution required analysis of higher-order peaks (J_03_, J_04_, and J_n_), the mean PF number was derived from a second fitting of higher-order peaks. For this, we assumed a PF spacing of 5.7 nm^56^ and considered electron microscopy observations that indicate a Gaussian distribution of the PF numbers^57^, and hence, we modeled a Gaussian centered on the mean. The model was able to reconstruct the scattering profile of MT solutions as a linear combination of the calculated profiles of individual species, suggesting additional information that could be obtained from full radial integration. Meridional peaks were fitted using a four-parameter Lorentzian function to determine the axial repeat distance by fitting the 1 nm⁻¹ layer line, I(q) = y₀ + a / [1 + ((q – q₀) / b)²].

#### Synthesis of compounds 1a and 1b (NMR spectra)

Unless otherwise stated reagents were purchased from general suppliers (Merck, Fluorochem, TCI) and used wCDClithout further purification. PTX was kindly supplied by Indena s.p.a. (Italy). All solvents were of reagent grade or HPLC grade. All reactions were carried out in oven-dried glassware and dry solvents, under nitrogen atmosphere, and were monitored by glasses or aluminum TLC on silica gel (Merck precoated 60F254 plates), with detection by UV light (254 nm), or by TLC stains as potassium permanganate, bromocresol green or panisaldehyde stain. Purification of intermediates and final products was mostly carried out by flash column chromatography, using high purity grade silicagel (Merck Grade, pore size 60 Å, 230-400 mesh particle size, Sigma Aldrich) as stationary phase. Alternatively, purification was performed using Biotage Isolera® One system in direct pase using Biotage® Sfär Silica D cartridges (4/10/35 g). ^1^H-NMR and ^13^C-NMR spectra were recorded on a Bruker Avance Spectrometer 400 MHz using commercially available deuterated solvents (chloroform-d, dimethylsulfoxide-d6) at RT. Chemical shifts (δ) are reported in parts per million (ppm) and are reported relative to tetramethylsilane, used as an internal standard. Mass spectrometry (MS) analyses were performed by direct infusion of the samples into an LCQ Fleet ion trap mass spectrometer equipped with an electrospray ionization (ESI) source. MS-SCAN conditions: 50-2000 amu, SCAN-time: 0.2 sec. Specific optical rotation, or [α]D, was measured on a Jasco P-1030 polarimeter in chloroform. This intrinsic property of chiral compounds is defined by the formula [α]λ = α/γl, where α is the angle through which plane polarized light is rotated by a compound’s solution of mass concentration γ and path length l (in this case l is equal to 10 cm, the length of the polarimeter cell). The superscript indicates the Celsius temperature at which the measurement is carried out, while the wavelength of the measurement is reported in the subscript. The letter D indicates that the wavelength used corresponds to the sodium D line (589 nm). The standard deviation is calculated over 10 measurements.

##### 2’-carboxybenzoylpaclitaxel (7)

Benzyl chloroformate (990 μL, 7.02 mmol) was added in 10 eq. aliquots (165 μL per aliquot) at 10 min intervals to a solution of PTX (100 mg, 0.117 mmol) and dry pyridine (950 μL, 11.71 mmol) in dry CH_2_Cl_2_ (5 mL), at RT and under N_2_ atmosphere. The reaction was stirred for 24 h, after which it was quenched with 2 mL of saturated aqueous NH_4_Cl. After the addition of AcOEt (10 mL), the organic phase was washed with saturated aqueous NH_4_Cl (3 x 5 mL) and brine (2 x 5 mL). The organic layer was then dried over Na_2_SO_4_, and the solvent was evaporated under reduced pressure. The crude product was purified by flash chromatography (silicagel, eluent mixture: 1:1 n-hexane/AcOEt) to obtain pure 7 (80.8 mg, 0.082 mmol, 70% yield).

^1^H NMR (400 MHz, CDCl_3_) δ 8.17 (d, *J* = 7.4 Hz, 2H, o-2-BzO), 7.76 (d, *J* = 7.1 Hz, 2H, o-3’-NHBz), 7.68 – 7.59 (m, 1H, p-2-BzO), 7.56-7.50 (m, 3H, m-2-BzO, p-3’-NHBz), 7.49 – 7.34 (m, 12H, m-3’-NHBz, o,m,p-2’-CbzO, o,m,p-3’-Ph), 6.98 (d, *J* = 9.3 Hz, 1H, 3’-NH), 6.37 – 6.28 (m, 2H, 10-H, 13-H), 6.02 (dd, *J* = 9.3, 2.7 Hz, 1H, 3’-H), 5.72 (d, *J* = 7.1 Hz, 1H, 2-H), 5.48 (d, *J* = 2.7 Hz, 1H, 2’-H), 5.26 – 5.14 (m, 2H, 4’-H), 5.01 (dd, *J* = 9.7, 2.3 Hz, 1H, 5-H), 4.47 (dd, *J* = 10.9, 6.6 Hz, 1H, 7-H), 4.35 (d, *J* = 8.5 Hz, 1H, 20a-H), 4.24 (d, *J* = 8.5 Hz, 1H, 20b-H), 3.85 (d, *J* = 7.0 Hz, 1H, 3-H), 2.59 (ddd, *J* = 14.7, 9.7, 6.5 Hz, 1H, 6a-H), 2.49 (s, 3H, 4-OAc), 2.47 – 2.41 (m, 1H, 14a-H), 2.26 (s, 3H, 10-OAc), 2.28 – 2.22 (m, 3H, 14a-H), 1.96 (s, 3H, 18-H), 1.98 – 1.86 (m, 1H, 6b-H), 1.72 (s, 3H, 19-H), 1.28 (s, 3H, 17-H), 1.17 (s, 3H, 16-H).

^13^C NMR (101 MHz, CDCl_3_) δ 203.97, 171.42, 170.01, 168.01, 167.23, 167.20, 142.89, 133.84, 132.97, 132.17, 130.39, 129.26, 129.06, 128.90, 128.85, 128.66, 128.56, 127.31, 126.74, 84.61, 81.24, 79.39, 77.02, 76.62, 75.76, 75.28, 72.31, 72.23, 70.93, 58.70, 52.91, 45.72, 35.76, 35.68, 27.00, 22.85, 22.32, 20.98, 14.96, 9.76 (detectable peaks).

##### Hexanedioic acid mono(2-(trimethylsilyl)ethyl) ester (5a)

Hexanedioic acid (1 g, 6.84 mmol) was added to a mixture of dry CH_2_Cl_2_ and dry pyridine (25 and 2.5 mL respectively, 10:1 ratio) and stirred at RT, under N_2_ atmosphere, until complete dissolution. Then 2-(trimethylsilyl) ethanol (392 μL, 2.74 mmol), EDC·HCl (788 mg, 4.11 mmol) and DMAP (167 mg, 1.39 mmol) were added, and the solution was stirred for 24 h. The reaction mixture was washed with phosphoric acid 10% (2 x 15 mL) and brine (20 mL). The organic phase was then dried over Na_2_SO_4_, and the solvent was evaporated under reduced pressure. The mono-protected product was isolated by flash chromatography (silicagel, eluent mixture: 8:2 n-hexane/AcOEt + 1% formic acid) to obtain pure 5a (446 mg, 1.8 1 mmol, 66% yield).

^1^H NMR (400 MHz, CDCl_3_) δ 10.94 (br s, 1H, COOH), 4.09 – 3.94 (m, 2H, 5-H), 2.29 – 2.06 (m, 4H, 1-H, 4-H), 1.54 (br s, 4H, 2-H, 3-H), 0. 91 – 0.76 (m, 2H, 6-H),-0.12 (m, 9H, SiMe3).

^13^C NMR (101 MHz, CDCl_3_) δ 178.87, 173.45, 62.43, 33.83, 33.42, 24.08, 23.89, 17.08,-1.82.

##### Nonanedioic acid mono(2-(trimethylsilyl)ethyl) ester (5b)

Nonanedioic acid (1 g, 5.31 mmol) was added to a mixture of dry CH_2_Cl_2_ and dry pyridine (25 and 2.5 mL respectively, 10:1 ratio) and stirred at RT, under N_2_ atmosphere, until complete dissolution. Then 2-(trimethylsilyl) ethanol (304 μL, 2.12 mmol), EDC·HCl (612 mg, 3.19 mmol) and DMAP (130 mg, 1.06 mmol) were added, and the solution was stirred for 24 h. The reaction mixture was washed with phosphoric acid 10% (2 x 15 mL) and brine (20 mL). The organic phase was then dried over Na_2_SO_4_, and the solvent was evaporated under reduced pressure. The mono-protected product was isolated by flash chromatography (silicagel, eluent mixture: 75:25 n-hexane/AcOEt + 1% formic acid) to obtain pure 5b (430mg, 1.49 mmol, 70% yield).

^1^H NMR (400 MHz, CDCl_3_) δ 10.98 (s, 1H, COOH), 4.15 – 4.05 (m, 2H, 8-H), 2.33 – 2.16 (m, 4H, 1-H, 7-H), 1.63 – 1.49 (m, 4H, 2-H, 6-H), 1.32 – 1.21 (m, 6H, 3-H, 4-H, 5-H), 0.97 – 0.86 (m, 2H, 9-H), 0.01 (s, 9H, SiMe3).

^13^C NMR (101 MHz, CDCl_3_) δ 179.99, 174.13, 62.51, 34.52, 34.09, 28.97, 28.93, 28.91, 24.94, 24.66, 17.39,-1.43.

##### 2’-O-Cbz-7-O-(1’’-O-(2’’’-(trimethylsilyl)ethyl)hexanedioyl)-paclitaxel (4a)

A solution of acid 5a (49 mg, 0.2 mmol) in CH_2_Cl_2_ (1 mL) was added to a stirred solution of 2’-O-Cbz-paclitaxel 7 (50 mg, 0.05 mmol) and DMAP (6 mg, 0.05 mmol) in CH_2_Cl_2_ (1 mL), at RT and under N_2_ atmosphere. Then DCC (42 mg, 0.2 mmol) was added, and the reaction mixture was stirred at RT for 18 h. The mixture was diluted with CH_2_Cl_2_ (5 mL) and it was filtered to remove the urea formed as a byproduct. The collected organic phase was washed with saturated aqueous NH_4_Cl (2 x 5 mL) and brine (2 x 5 mL), then it was dried over Na_2_SO_4_, and the solvent was evaporated under reduced pressure. The resulting crude was purified by flash chromatography (silicagel, eluent mixture: 6.5:3.5 n-hexane/AcOEt) to obtain compound 4a (55 mg, 0.045 mmol, 90% yield).

^1^H NMR (400 MHz, CDCl_3_) δ 8.p (d, 2H, o-2-BzO), 7.62 – 7.55 (m, 2H, o-3’-NHBz), 7.51 – 7.42 (m, 1H, p-2-BzO), 7.41 – 7.30 (m, 3H, m-2-BzO, p-3’-NHBz), 7.30 – 7.16 (m, 12H, m-3’-NHBz, o,m,p-2’-CbzO, o,m,p-3’-Ph), 6.78 (d, *J* = 9.3 Hz, 1H, 3’-NH), 6.13 (s, 1H, H-10), 6.16 – 6.07 (m, 1H, H-13), 5.84 (dd, *J* = 9.3, 2.7 Hz, 1H, 3’-H), 5.55 (d, *J* = 6.9 Hz, 1H, 2-H), 5.45 (dd, *J* = 10.5, 7.0 Hz, 1H, 7-H), 5.32 (d, *J* = 2.7 Hz, 1H, 2’-H), 5.08 – 4.96 (m, 2H, 4’-H), 4.82 (dd, *J* = 9.6, 1.9 Hz, 1H, 5-H), 4.19 (d, *J* = 8.4 Hz, 1H, 20a-H), 4.05 (d, *J* = 8.4 Hz, 1H, 20b-H), 4.04 – 3.98 (m, 3H, 5’’-H), 3.81 (d, *J* = 6.9 Hz, 1H, 3-H), 2.46 (ddd, *J* = 14.4, 9.6, 7.1 Hz, 1H, 6a-H), 2.30 (s, 3H, 4-OAc), 2.25 (m, 2H, 14a-H), 2.19 – 2.10 (m, 4H, 1’’-H, 4’’-H), 2.07 (m, 14b-H), 2.01 (s, 3H, 10-OAc), 1.85 (s, 3H, 18-H), 1.67 (s, 3H, 19-H), 1.73 – 1.67 (m, 1H, 6b-H) 1.54 – 1.44 (m, 4H, 2’’-H, 3’’-H), 1.07 (s, 3H, 17-H), 1.02 (s, 3H,16-H), 0.89 – 0.79 (m, 2H, 6’’-H),-0.11 (s, 9H, SiMe3).

^13^C NMR (101 MHz, CDCl_3_) δ 201.87, 173.45, 172.17, 169.38, 168.60, 167.77, 167.08, 166.72, 153.82, 140.90, 136.50, 134.12, 133.50, 133.34, 132.36, 131.81, 130.01, 128.97, 128.90, 128.68, 128.54, 128.52, 128.49, 128.34, 128.29, 126.96, 126.40, 83.81, 80.69, 78.46, 76.55, 76.17, 75.02, 74.39, 71.83, 71.02, 70.61, 62.25, 55.85, 52.59, 46.68, 43.09, 35.21, 33.97, 33.45, 33.17, 26.25, 24.13, 23.72, 22.44, 21.09, 20.51, 17.11, 14.31, 10.69,-1.69.

##### Synthesis of 6-O-(2’,3’-dihydroinden-1’-yl)-1-O-(2’’-(trimethylsilyl)ethyl) hexanedioate (10a)

DMAP (26 mg, 0.21 mmol), EDC·HCl (119 mg, 0.62 mmol), and 1-indanol (87 mg, 0.62 mmol) were added sequentially to a solution of 5a (101.5 mg, 0.41 mmol) in dry CH_2_Cl_2_ and pyridine (18 and 2 mL respectively, 10:1 ratio), at RT and under N_2_ atmosphere. The reaction mixture was stirred for 6 h. When TLC monitoring confirmed reaction completion, HCl 1M was added (40 mL). The aqueous phase was extracted with CH_2_Cl_2_ (4 x 15 mL), then the collected organic layer was dried over Na_2_SO_4_, and the solvent was evaporated under reduced pressure. The crude oil was purified by Biotage® direct phase flash chromatography (eluent mixture: n-hexane/AcOEt from 98:2 to 80:20) to afford pure compound 10a (90.9 mg, 0.25 mmol, 61%).

^1^H NMR (400 MHz, CDCl_3_) δ 7.39 (d, *J* = 7.4 Hz, 1H, 10-H), 7.31 – 7.16 (m, 3H, 13-H, 12-H, 11-H), 6.21 (dd, *J* = 6.9, 3.9 Hz, 1H, 7-H), 4.20 – 4.11 (m, 2H, H-2), 3.16 – 3.04 (m, 1H, 9a-H), 2.92 – 2.81 (m, 1H, 9b-H), 2.56 – 2.43 (m, 1H, 8a-H), 2.36 – 2.24 (m, 4H, 3-H, 6-H), 2.13 – 2.01 (m, 1H, 8b-H), 1.71 – 1.61 (m, 4H, 4-H, 5-H), 1.02 – 0.93 (m, 2H, 1-H), 0.04 (s, 9H,-SiMe3).

^13^C NMR (101 MHz, CDCl_3_) δ 173.45, 173.35, 144.39, 141.19, 128.95, 126.76, 125.55, 124.85, 78.31, 62.53, 34.22, 34.14, 32.40, 30.26, 24.55, 24.47, 17.41,-1.42.

##### Hexanedioic acid mono(2’,3’-dihydroinden-1’-yl) ester (9a)

TBAF 1M in THF (1 mL, 1 mmol) was added dropwise to a stirred solution of 10a (90 mg, 0.25 mmol) in dry THF (10 mL), at – 5 °C and under N_2_ atmosphere. The reaction mixture was warmed up to RT and stirred for 4 h. When TLC monitoring confirmed reaction completion, saturated aqueous NH_4_Cl (15 mL) was added. The aqueous phase was extracted with AcOEt (4 x 15 mL), then the collected organic layer was dried over Na_2_SO_4_ and evaporated under reduced pressure. The crude oil was purified by flash chromatography (silicagel, eluent mixture: 8:2 n-hexane/AcOEt + 1% formic acid) to obtain pure 9a (41.3 mg, 0.157 mmol, 63% yield).

^1^H NMR (400 MHz, CDCl_3_) δ 7.38 (d, *J* = 7.4 Hz, 1H, 8-H), 7.37 – 7.05 (m, 3H, 11-H, 10-H, 9-H), 6.20 (dd, *J* = 6.9, 3.8 Hz, 1H, 5-H), 3.15 – 3.03 (m, 1H, 7a-H), 2.92 – 2.80 (m, 1H, 7b-H), 2.55 – 2.43 (m, 1H, 6a-H), 2.39 – 2.29 (m, 4H, 1-H, 4-H), 2.18 – 2.00 (m, 1H, 6b-H), 1.79 – 1.54 (m, 4H, 2-H, 3-H).

^13^C NMR (101 MHz, CDCl_3_) δ 173.42, 144.50, 141.18, 129.05, 126.84, 125.62, 124.93,

78.46, 34.23, 33.52, 32.44, 30.32, 24.46, 24.19.

##### 2’-O-Cbz-7-O-(6’’-O-(2’’’,3’’’-dihydroinden-1’’’-yl)hexanedioyl)paclitaxel (2a)

A solution of compound 9a (28 mg, 0.107 mmol) in CH_2_Cl_2_ (2 mL) was added to a stirred solution of 2’-O-Cbzpaclitaxel 7 (70 mg, 0.071 mmol) and DMAP (8.7 mg, 0.071 mmol) in CH_2_Cl_2_ (3 mL), at RT and under N_2_ atmosphere. Then, DCC (58 mg, 0.284 mmol) was added and the reaction mixture was stirred at RT for 29 h. The mixture was diluted with CH_2_Cl_2_ (8 mL) and filtered on a plug of celite. The organic phase was washed with saturated aqueous NH_4_Cl (2 x 10 mL) and brine (2 x 10 mL), then it was dried over Na_2_SO_4_, and the solvent was evaporated under reduced pressure. The resulting crude was purified by flash chromatography (silicagel, eluent mixture: 7:3 n-hexane/AcOEt) to obtain compound 2a (61.1 mg, 0.050 mmol, 70% yield).

^1^H NMR (400 MHz, CDCl_3_) δ 8.12 (d, *J* = 7.3 Hz, 2H, o-2-BzO), 7.73 (d, *J* = 7.3 Hz, 2H, o-3’-NHBz), 7.64 – 7.56 (m, 1H p-2-BzO), 7.54 – 7.44 (m, 3H, m-2-BzO, p-3’-NHBz), 7.43 – 7.29 (m, 13H, m-3’-NHBz, o,m,p-2’-CbzO, o,m,p-3’-Ph, 8’’-H), 7.29 – 7.18 (m, 3H, 9’’-H, 10’’-H, 11’’-H), 6.95 (d, *J* = 9.3 Hz, 1H, 3’-NH), 6.28 – 6.20 (m, 2H, 10-H, 13-H), 6.19 (dd, *J* = 7.0, 3.9 Hz, 1H, 5’’-H), 5.97 (dd, *J* = 9.3, 2.5 Hz, 1H, 3’-H), 5.69 (d, *J* = 6.9 Hz, 1H, 2-H), 5.58 (dd, *J* = 10.4, 7.1 Hz, 1H, 7-H), 5.46 (d, *J* = 2.7 Hz, 1H, 2’-H), 5.21 – 5.10 (m, 2H, 4’-H), 4.95 (d, *J* = 8.9 Hz, 1H, 5-H), 4.32 (d, *J* = 8.4 Hz, 1H, 20a-H), 4.19 (d, *J* = 8.4 Hz, 1H, 20b-H), 3.95 (d, *J* = 6.9 Hz, 1H, 3-H), 3.15 – 3.03 (m, 1H, 7’’a-H), 2.86 (ddd, *J* = 16.0, 8.5, 5.1 Hz, 1H, 7’’b-H), 2.59 (ddd, *J* = 14.5, 9.4, 7.4 Hz, 1H, 6a-H), 2.56 – 2.44 (m, 1H, 6’’a-H), 2.44 (s, 3H, 4-OAc), 2.41 – 2.16 (m, 6H, 14-H, 1’’-H, 4’’-H), 2.12 (s, 3H, 10-OAc), 2.11 – 2.00 (m, 2H), 1.99 (s, 3H, 18-H), 1.94 – 1.82 (m, 1H, 6b-H), 1.80 (s, 3H, 19-H), 1.71 – 1.57 (m, 4H, 2’’-H, 3’’-H), 1.20 (s, 3H, 17-H), 1.16 (s, 3H, 16-H).

^13^C NMR (101 MHz, CDCl_3_) δ 202.06, 172.34, 169.59, 168.79, 167.98, 166.99, 141.14, 136.78, 133.72, 133.60, 132.57, 132.00, 130.23, 129.11, 128.90, 128.86, 128.75, 128.71, 128.55, 128.50, 127.17, 126.72, 126.63, 125.53, 124.78, 84.03, 80.92, 78.75, 78.19, 76.81, 76.41, 75.24, 74.62, 72.04, 71.23, 70.82, 56.09, 52.84, 46.91, 43.31, 35.45, 34.23, 33.95, 33.65, 32.36, 30.21, 26.48, 24.39, 23.90, 22.66, 21.30, 20.70, 14.53, 10.90 (detectable peaks).

##### 7-O-(6’’-O-(2’’’,3’’’-dihydroinden-1’’’-yl)hexanedioyl) paclitaxel (1a)

Pd/C (10% w/w, 3 mg) was added to a solution of 2a (30 mg, 0.024 mmol) in dry MeOH (2 mL) under hydrogen atmosphere. The reaction mixture was stirred at RT, and it was monitored by TLC until completion (5 h). The reaction mixture was then filtered on a celite pad, which was washed with MeOH (5 mL x2). The organic phase was dried with Na_2_SO_4_, and the solvent was evaporated under reduced pressure to afford pure compound 1a in high yield (27.6 mg, 0.0238 mmol, 98% yield).

^1^H NMR (400 MHz, CDCl_3_) δ 8.13 (d, *J* = 7.3 Hz, 2H, o-2-BzO), 7.79 (d, *J* = 7.3 Hz, 2H, o-3’-NHBz), 7.68 – 7.59 (m, 1H, p-2-BzO), 7.57 – 7.33 (m, 11H, m-2-BzO, m,p-3’-NHBz, o,m,p-3’-Ph, 8’’-H), 7.33 – 7.19 (m, 4H, 3’-NH, 9’’-H, 10’’-H, 11’’-H), 6.26 – 6.15 (m, 3H, 10-H, 13-H, 5’’-H), 5.87 – 5.80 (m, 1H, 3’-H), 5.69 (d, *J* = 6.9 Hz, 1H, 2-H), 5.57 (dd, *J* = 10.3, 7.3 Hz, 1H, 7-H), 4.95 (d, *J* = 9.0 Hz, 1H, 5-H), 4.82 (d, *J* = 2.4 Hz, 1H, 2’-H), 4.33 (d, *J* = 8.4 Hz, 1H, 20a-H), 4.21 (d, *J* = 8.4 Hz, 1H, 20b-H), 3.93 (d, *J* = 6.7 Hz, 1H, 3-H), 3.18 – 3.06 (m, 1H, 7’’a-H), 2.89 (ddd, *J* = 15.9, 8.5, 5.1 Hz, 1H, 7’’b-H), 2.65 – 2.45 (m, 2H, 6a-H, 6’’a-H), 2.40 (s, 3H, 4-OAc), 2.38 – 2.20 (m, 6H, 14-H, 1’’-H, 4’’-H), 2.16 (s, 3H, 10-OAc), 2.16 – 2.04 (m, 1H, 6’’b-H), 1.88 – 1.79 (m, 7H, 6b-H, 18-H, 19-H), 1.75 – 1.57 (m, 4H, 2’’-H, 3’’-H), 1.21 (s, 3H, 17-H), 1.18 (s, 3H, 16-H).

^13^C NMR (101 MHz, CDCl_3_) δ 202.56, 174.14, 173.19, 173.13, 171.00, 169.57, 169.52, 167.69, 167.54, 157.68, 145.01, 141.80, 141.06, 138.81, 134.43, 133.61, 132.54, 130.84, 129.77, 129.59, 129.53, 129.39, 129.33, 128.89, 127.76, 127.37, 126.16, 125.45, 84.58, 81.72, 79.15, 78.86, 77.11, 75.92, 75.02, 74.04, 72.69, 71.88, 56.84, 55.63, 47.63, 43.90, 36.24, 34.86, 34.52, 34.31, 33.00, 30.86, 27.18, 25.03, 24.55, 23.22, 21.48, 21.38, 15.29, 11.48 (detectable peaks).

MS (ESI): m/z [M + Na]+ calcd. for C62H67NO17Na: 1120.43, found: 1120.75. [α]_D_^25^ ± SD:-290°± 6.

##### 2’-(trimethylsilyl)ethyl (S)-9-((7’’-methoxy-1’’,2’’,3’’,4’’-tetrahydro naphthalen-2’’-yl)amino)-9-oxononanoate (10b)

DMAP (29.6 mg, 0.243 mmol), EDC·HCl (99.8 mg, 0.52 mmol) and (S)-7-methoxy-2-aminotetralin (92.2 mg, 0.52 mmol) were added sequentially to a solution of 5b (99.5 mg, 0.345 mmol) in dry CH_2_Cl_2_ and pyridine (15 and 1.5 mL respectively, 10:1 ratio), at RT and under N_2_ atmosphere. The reaction mixture was stirred for 4 h. When TLC monitoring confirmed reaction completion, HCl 1M was added (35 mL). The aqueous phase was extracted with CH_2_Cl_2_ (3 x 15 mL), then the collected organic layer was dried over Na_2_SO_4_, and the solvent was evaporated under reduced pressure. The crude oil was purified by direct phase flash chromatography (silicagel, eluent mixture: 85:25 n-hexane/AcOEt + 1% formic acid) to afford pure compound 10b in high yield (153.8 mg, 0.344 mmol, 98%).

^1^H NMR (400 MHz, CDCl_3_) δ 6.96 (d, *J* = 8.4 Hz, 1H, 13-H), 6.67 (dd, *J* = 8.4, 2.7 Hz, 1H, 14-H), 6.54 (d, *J* = 2.7 Hz, 1H, 15-H), 6.32 (d, *J* = 7.9 Hz, 1H, 10-NH), 4.28 – 4.15 (m, 1H, 10-H), 4.16 – 4.10 (m, 2H, 2-H), 3.72 (s, 3H, 14-OMe), 3.03 (dd, *J* = 16.4, 5.3 Hz, 1H, 16a-H), 2.86 – 2.68 (m, 2H, 12-H), 2.61 (dd, *J* = 16.4, 8.5 Hz, 1H, 16b-H), 2.24 (t, *J* = 7.5 Hz, 2H, 3-H), 2.20 – 2.12 (m, 2H, 9-H), 2.03 – 1.93 (m, 1H, 11a-H), 1.72 (dtd, *J* = 12.6, 9.1, 6.2 Hz, 1H, 11b-H), 1.65 – 1.51 (m, 4H, 4-H, 8-H), 1.28 (br s, 6H, 5-H, 6-H, 5-H), 1.01 – 0.92 (m, 2H, 1-H), 0.03 (s, 9H, SiMe).

^13^C NMR (101 MHz, CDCl_3_) δ 174.59, 173.26, 135.89, 130.37, 128.22, 114.61, 113.20, 63.03, 55.89, 55.88, 45.54, 37.52, 36.63, 35.10, 29.69, 29.61, 29.55, 29.47, 26.83, 26.36, 25.53, 17.99,-1.43

##### (S)-9-((7’-methoxy-1’,2’,3’,4’-tetrahydronaphthalen-2’-yl)amino)-9-oxononanoic acid (9b)

TBAF 1M in THF (1.117 mL, 1.117 mmol) was added dropwise to a stirred solution of 10b (125 mg, 0.279 mmol) in dry THF (11 mL), at – 5 °C and under N_2_ atmosphere. The reaction mixture was then warmed up to RT and stirred for 4 h. When TLC monitoring confirmed reaction completion, saturated aqueous NH_4_Cl (15 mL) was added. The aqueous phase was extracted with AcOEt (3 x 15 mL), then the collected organic layer was dried over Na_2_SO_4_, and the solvent was evaporated under reduced pressure. The crude oil was purified by Biotage® direct phase flash chromatography (eluent mixture: n-hexane/AcOEt + 1% formic acid from 90:10 to 20:80) to obtain pure 9b (84.6 mg, 0.243 mmol, 87% yield).

^1^H NMR (400 MHz, CDCl_3_) δ 7.01 (d, *J* = 8.4 Hz, 1H, 11-H), 6.71 (dd, *J* = 8.4, 2.7 Hz, 1H, 12-H), 6.59 (d, *J* = 2.7 Hz, 1H, 13-H), 5.49 (d, *J* = 7.8 Hz, 1H, 8-NH), 4.37 – 4.24 (m, 1H, 8-H), 3.77 (s, 3H, OMe), 3.09 (dd, *J* = 16.4, 5.1 Hz, 1H, 14a-H), 2.90 – 2.71 (m, 2H, 10-H), 2.61 (dd, *J* = 16.4, 7.6 Hz, 1H, 14b-H), 2.34 (t, *J* = 7.5 Hz, 2H, 1-H), 2.20 – 2.11 (m, 2H, 7-H), 2.07 – 1.95 (m, 1H, 9a-H), 1.84 – 1.71 (m, 1H, 9b-H), 1.68 – 1.56 (m, 4H, 2-H, 6-H), 1.32 (br s, 6H, 3-H, 4-H, 5-H).

^13^C NMR (101 MHz, CDCl_3_) δ 179.37, 173.75, 158.41, 135.81, 130.40, 128.21, 114.63, 113.25, 55.92, 45.67, 37.48, 36.54, 34.67, 29.63, 29.55, 29.50, 29.39, 26.77, 26.37, 25.32.

##### 2’-O-Cbz-7-O-((S)-9’’-(7’’’-methoxy-1’’’,2’’’,3’’’,4’’’-tetrahydronaphthalen-2’’’-yl)amino)-9’’-oxononanoyl) paclitaxel (2b)

A solution of compound 9b (30.6 mg, 0.088 mmol) in CH_2_Cl_2_ (1.5 mL) was added to a stirred solution of 2’-OCbz-paclitaxel 7 (58 mg, 0.059 mmol) in CH_2_Cl_2_ (1.5 mL), at RT and under N_2_ atmosphere. Subsequently, DCC (49 mg, 0.236 mmol) and DMAP (10 mg, 0.082 mmol) were added and the reaction mixture was stirred at RT for 26 h. The mixture was diluted with CH_2_Cl_2_ (10 mL) and it was filtered on a plug of celite. The organic layer was washed with saturated aqueous NH_4_Cl (2 x 10 mL) and brine (2 x 10 mL), then it was dried over Na_2_SO_4_, and the solvent was evaporated under reduced pressure. The resulting crude was purified by Biotage® direct phase flash chromatography (eluent mixture: n-hexane/AcOEt from 88:12 to 0:100) to obtain compound 2a (67.5 mg, 0.051 mmol, 87% yield).

^1^H NMR (400 MHz, CDCl_3_) δ 8.12 (d, *J* = 7.2 Hz, 2H, o-2-BzO), 7.73 (d, *J* = 7.2 Hz, 2H, o-3’-NHBz), 7.64 – 7.56 (m, 1H, p-2-BzO), 7.54 – 7.44 (m, 3H, m-2-BzO, p-3’-NHBz), 7.42 – 7.28 (m, 12H, m-3’-NHBz, o,m,p-2’-CbzO, o,m,p-3’-Ph), 7.04 (d, *J* = 9.3 Hz, 1H, 3’-NH), 6.99 (d, *J* = 8.6 Hz, 1H, 11’’-H), 6.70 (d, *J* = 8.5 Hz, 1H, 12’’-H), 6.58 (d, *J* = 2.9 Hz, 1H, 13’’-H), 6.31 – 6.20 (m, 2H, 10-H, 13-H), 5.97 (dd, *J* = 9.3, 2.9 Hz, 1H, 3’-H), 5.68 (dd, *J* = 16.5, 7.5 Hz, 2H, 2-H, 7-H), 5.64 – 5.54 (m, 1H, 8’’-NH), 5.46 (d, *J* = 2.8 Hz, 1H, 2’-H), 5.21 – 5.10 (m, 2H, 4’-H), 4.95 (d, *J* = 9.3 Hz, 1H, 5-H), 4.32 (d, *J* = 8.4 Hz, 1H, 20a-H), 4.29 – 4.22 (m, 1H, 8’-H), 4.19 (d, *J* = 8.4 Hz, 1H, 20b-H), 3.95 (d, *J* = 6.9 Hz, 1H, 3-H), 3.74 (s, 3H, OMe), 3.07 (dd, *J* = 16.4, 5.1 Hz, 1H, 14’’a-H), 2.86 – 2.72 (m, 2H, 10’’-H), 2.60 (dd, *J* = 16.6, 7.8 Hz, 2H, 14’’b-H, 6a-H), 2.44 (s, 3H, 4-OAc), 2.42 – 2.17 (m, 4H, 14a-H, 1’’-H), 2.16 – 2.09 (m, 5H, 10-OAc, 7’’-H), 2.00 (s, 3H, 18-H), 1.95 – 1.84 (m, 2H, 6b-H, 9’’a-H), 1.81 (s, 3H, 19-H), 1.77 – 1.64 (m, 1H, 9’’b-H), 1.63 – 1.52 (m, 4H, 2’’-H, 8’’-H), 1.34 – 1.23 (m, 6H, 3’’-H, 4’’-H, 5’’-H), 1.21 (s, 3H, 17-H), 1.16 (s, 3H, 16-H).

^13^C NMR (101 MHz, DMSO-d6) δ 201.59, 171.86, 171.58, 169.87, 169.12, 168.65, 166.33, 165.21, 157.21, 156.61, 153.87, 139.81, 137.01, 135.94, 134.92, 134.07, 133.60, 132.70, 131.57, 129.77, 129.59, 129.35, 128.80, 128.71, 128.55, 128.36, 128.29, 127.57, 127.41, 127.37, 113.50, 112.15, 82.81, 79.75, 77.15, 76.60, 74.50, 74.05, 71.01, 70.83, 69.81, 55.28, 54.91, 54.08, 47.51, 44.57, 42.91, 35.40, 35.33, 33.33, 31.69, 28.92, 28.49, 28.42, 26.61, 26.20, 25.31, 24.44, 22.51, 21.17, 20.47, 13.92, 10.59 (detectable peaks).

##### 7-O-((S)-9’’-(7’’’-methoxy-1’’’,2’’’,3’’’,4’’’-tetrahydronaphthalen-2’’’-yl)amino) - 9’’-oxononanoyl) paclitaxel (1b)

Pd/C (10% w/w, 4.6 mg) was added to a solution of 2b (57 mg, 0.043 mmol) in dry MeOH (4.25 mL) under hydrogen atmosphere. The reaction mixture was stirred at RT, and it was monitored by TLC until completion (4 h). The reaction mixture was then filtered on a celite pad, which was washed with MeOH (5 mL x2). The organic phase was dried with Na_2_SO_4_, and the solvent was evaporated under reduced pressure. The crude product was purified by Biotage® direct phase flash chromatography (eluent mixture: CH_2_Cl_2_/MeOH from 100:0 to 96:4) to afford pure compound 1b (27.6 mg, 0.0238 mmol, 45% yield).

^1^H NMR (400 MHz, CDCl_3_) δ 8.12 (d, *J* = 7.1 Hz, 2H, o-2-BzO), 7.78 (d, *J* = 7.1 Hz, 2H, o-3’-NHBz), 7.66 – 7.58 (m, 1H, p-2-BzO), 7.55 – 7.31 (m, 10H, m-2-BzO, m,p-3’-NHBz, o,m,p-3’-Ph), 7.24 (d, *J* = 9.0 Hz, 1H, 3’-NH), 7.01 (d, *J* = 8.4 Hz, 1H, 11’’-H), 6.71 (dd, *J* = 8.4, 2.9 Hz, 1H, 12’’-H), 6.60 (d, *J* = 2.8 Hz, 1H, 13’’-H), 6.24 (s, 1H, 10-H), 6.17 (t, *J* = 8.2 Hz, 1H, 13-H), 5.81 (dd, *J* = 9.0, 2.7 Hz, 1H, 3’-H), 5.67 (d, *J* = 6.9 Hz, 1H, 2-H), 5.55 (dd, *J* = 10.6, 7.2 Hz, 2H, 7-H, 8’’-NH), 4.94 (d, *J* = 7.6 Hz, 1H, 5-H), 4.79 (d, *J* = 2.6 Hz, 1H, 2’-H), 4.35 – 4.23 (m, 2H, 20a-H, 8’’-H), 4.19 (d, *J* = 8.5 Hz, 1H, 20b-H), 3.92 (d, *J* = 6.9 Hz, 1H, 3-H), 3.77 (d, *J* = 2.6 Hz, 3H, OMe), 3.09 (dd, *J* = 16.5, 5.3 Hz, 1H, 14’’a-H), 2.89 – 2.73 (m, 2H, 10’’-H), 2.66 – 2.53 (m, 2H, 14’’b-H, 6a-H), 2.38 (s, 3H, 4-OAc), 2.36 – 2.29 (m, 3H, 14a-H, 1’’-H), 2.29 – 2.17 (m, 1H, 14b-H), 2.15 (d, *J* = 6.9 Hz, 5H, 10-OAc, 7’’-Hp), 2.02 – 1.85 (m, 1H, 9’’a-H), 1.82 (d, J = 8.0 Hz, 7H, 18-H, 19-H, 6b-H), 1.80 – 1.65 (m, 1H, 9’’b-H), 1.59 (d, *J* = 13.3 Hz, 4H, 2’’-H, 8’’-H), 1.34 – 1.22 (m, 6H, 3’’-H, 4’’-H, 5’’-H), 1.20 (s, 3H, 17-H), 1.17 (s, 3H, 16-H).

^13^C NMR (101 MHz, CDCl_3_) δ 202.08, 173.04, 172.78, 172.53, 170.43, 168.96, 167.09, 167.02, 157.85, 140.52, 138.28, 135.37, 133.90, 133.09, 132.01, 130.30, 129.87, 129.23, 129.06, 128.86, 128.80, 128.37, 127.72, 127.22, 114.07, 112.71, 84.06, 81.17, 78.62, 76.59, 75.40, 74.47, 73.47, 72.15, 71.25, 56.34, 55.39, 55.08, 47.11, 45.01, 43.37, 37.07, 36.10, 35.71, 34.17, 33.99, 33.61, 29.16, 29.01, 28.93, 26.66, 25.71, 25.03, 24.51,22.67, 20.92, 20.88, 14.76, 10.96.

MS (ESI): m/z [M + Na]+ calcd. for C67H78N2O17Na: 1205.52, found: 1205.44. [α]_D_^25^ ± SD:-235°± 5.

#### Cell Biology

Human NSCLC carcinoma-derived cells A549, human cervical carcinoma HeLa, HeLaβIII (MDR overexpressing βIII tubulin isotype), Kb-3.1 (HeLa subclone), Kb-V1 (MDR overexpressing PgP) and human SH-SY5Y neuroblastoma cell lines were cultured at 37 °C in DMEM supplemented with 10% fetal calf serum, 2 mM L-glutamine, 1 mM sodium pyruvate, 40 μg/mL gentamycin, 100 IU/ml penicillin and 100 μg/mL streptomycin in a 5% CO_2_ air atmosphere.

Indirect immunofluorescence images were obtained using A549 cells plated at a density of 35,000 cells/well onto 18 mm round coverslips, cultured overnight, and treated with increasing amounts of compounds **1a**, **1b**, PTX, or the drug vehicle (DMSO) for 24 h. DMSO was always less than 0.5% v/v. PEG and Triton X-100 were applied to cells prior the fixation with 3.7% formaldehyde, as previously described^58^. Cells were incubated with a DM1A mouse monoclonal antibody (Sigma-Aldrich) reacting with α-tubulin. After that, samples were washed and incubated with AF488 goat anti-mouse polyclonal antibody (ThermoFisher), and 3 μM DAPI (Merk) was added for staining the DNA. Images were taken using a Leica DM 6000 B epifluorescence microscope employing a 100X objective with a N.A. of 1.46. Images of α-tubulin and DNA were usually taken at slightly different but close z planes in order to improve the clarity of the focus for each individual image. Images were recorded using a Leica DFC360 FX CCD camera.

Antiproliferative assays were performed as described^23^. Briefly, the MTT viability assay was used for the determination of the cytotoxicity of compounds **1a**, **1b** and PTX. A549, HeLa, HeLaβIII, Kb-3.1, and Kb-V cells were seeded at densities of 65,000 cells/well (A549, HeLa, Kb-3.1) or 100,000 cells/well (HeLaβIII, Kb-V1), and were grown for 24 h at 37 °C and 5% CO_2_. Then, increasing concentrations of drugs were added and incubated for 48 h prior to the reaction with the MTT solution. IC50 values were obtained by fitting the experimental data (absorbance at 570/690 nm) to a four-parameter logistic curve using SigmaPlot 14.5 software package (Systat Software, Inc., San Jose, CA, USA) and IC50 values were expressed as mean ± SEM from three independent experiments, each performed in duplicate.

##### Kinesin and dynein intracellular tracking in SH-SY5Y cells

Cy5-KBP and Cy5-DBP tracking peptides were synthesized at CIB-Margarita Salas (CSIC), following a published protocol^27^. Briefly, the peptide chains were assembled via solid phase peptide synthesis (SPPS) using fluorenylmethyloxycarbonyl (Fmoc) as N-terminal protecting group and, subsequently labelled with the organic dye cyanine 5 (Cy5) via a maleimide-thiol reaction between the N-terminal cysteine residues of the peptides and the commercially available Cy5-maleimide. The final peptides were purified via reverse phase HPLC, dissolved in DMSO to a final 2.5 mM concentration and stored in small aliquots at −20 °C. SH-SY5Y cells were seeded into μ-Slide 8-wells ibiTreat plates (ibidi®, flat polymer coverslip-bottom, tissue culture treated) at a concentration of 35,000 cells/well and incubated for 48 h at 37 °C and 5% CO_2_. Solutions of the tested compounds at the desired concentrations (1 μM for PTX and FTX2, 2.5 μM for **1a** and **1b**) with and without 2.5 μM Cy5-KBP or Cy5-DBP were prepared in Gibco™ DMEM without phenol red, previously supplemented with 10% v/v fetal calf serum, 2 mM L-glutamine, 1 mM sodium pyruvate, 40 μg/mL gentamycin, 100 IU/mL penicillin and 100 μg/mL streptomycin, and incubated 20 min at 37 °C and 5% CO_2_. After this time, the culture medium was replaced by the drug working solution containing either Cy5-KBP or Cy5-DBP, and the plate was incubated for an additional 15 min at 37 °C and 5% CO_2_. Then, the culture medium was replaced with drug working solution without labelled peptide and the cells were imaged using a confocal laser scanning microscope Leica TCS SP8 with a 63x oil immersion objective that included a humidified incubation chamber, a CO_2_ controller and a heating unit. Cy5 dye was excited at 635 nm and the fluorescence emission was collected at 645-690 nm. Images were recorded every 1.793 s for 3 min as 2-layer-z-stacks. For each condition, two to three different fields were imaged in two separate experiments.

Single-peptide trajectories were tracked with the TrackMate plugin from the Fiji package software (https://imagej.net/TrackMate). Particles with an estimated blob diameter of 1 – 1.5 μm were detected using the Laplacian of Gaussian (LoG) detector and linked with the Linear motion LAP tracker. Only spots that were at a maximum distance of 2 μm were linked. Trajectories with less than 6 spots were discarded since they were probably artifactual, and track mean displacement and track mean velocity were calculated and normalized to control cells. Data were further analyzed with MATLAB to perform a Mean Square Displacement (MSD) analysis to determine the mode of displacement of the peptides followed over time^59,60^. Briefly, for each trajectory the MSD function was calculated and its log-log representation fitted with a linear function (𝑙𝑜𝑔(𝑀𝑆𝐷(τ)) = 2 x 𝑙𝑜𝑔(𝜏) + 𝐶). MSD curves with an R^2^ coefficient < 0.8 were removed. The slope (α) of each MSD curve determines the motion type of each particle^61^ and particles were classified into: constrained particles (CM) α < 1, Brownian particles (BM) α = 1, and transported particles (TM) α > 1. Two independent experiments were performed using three biological replicates (N = 6). Statistical analyses of normalized track mean displacement and track mean velocity were performed using an unpaired t-test with Welch’s correction in GraphPad Prism® 8 software. Statistical significance was set at **** p < 0.0001, *** p < 0.001, ** p < 0.01, and * p < 0.05.

##### PC12 culture and Fluorescence Decay After Photoactivation (FDAP)

PC12 cells (originally obtained from J.A. Wagner, Harvard Medical School) were cultured in serum-DMEM (DMEM supplemented with 10% fetal bovine serum and antibiotics (100 U/ml penicillin and 100 µg/ml streptomycin)) at 37 °C with 10 % CO_2_ in a humidified incubator. For live cell imaging, cells were transfected with the respective pRc/CMV expression vector using Lipofectamine 2000 (Thermo-Fisher Scientific, USA) as previously described^62^. FDAP experiments were performed essentially as previously described^63^. Briefly, cells expressing PAGFP-tagged Tau441 were plated on 35-mm glass-bottom culture dishes (MatTek, USA), transfected, and neuronally differentiated by medium exchange to serum-reduced DMEM containing 100 ng/mL 7S mouse NGF. After 3 days, the medium was exchanged to serum-reduced DMEM without phenol red with NGF, and the respective compound (or DMSO for carrier control) were added at the desired concentration. After 20 h, live cell imaging was performed using a laser scanning microscope (Nikon Eclipse Ti2-) equipped with a LU-N4 laser unit with 488-nm and 405-nm lasers and a Fluor 60× ultraviolet-corrected objective lens (NA 1.4) and a C2+scanner enclosed in an incubation chamber at 37 °C and 5 % CO_2_. Photoactivation was performed with a 405-nm laser using the microscope software (NIS-Elements version AR 5.02.03). The following steps were performed in a photoactivation experiment. A transfected cell with a suitable process (minimum length 30 µm) was selected in the field of view (50 µm × 50µm) and a pre-activation image was saved at 488-nm excitation. The scan window was reduced to a 6-μm diameter activation spot (corresponding to 130 pixels), and photoactivation was performed in the centre of the neurite using the 405-nm blue diode (laser intensity 1.0%; corresponding to an output of 30 µW) with two scans at a pixel dwell time of 4.08 μs (corresponding to a total activation time at the spot of 1.1 ms). The scan window was enlarged to the initial size, and the first scan was performed after 1 s with the 488-nm laser at the lowest practical laser intensity. Subsequent images were acquired at a rate of 1 frame/s. 112 images with a resolution of 256 × 256 pixels were collected per activated neurite. To ensure that the imaging conditions did not affect cell viability and subsequent evaluation, neurites were excluded from analysis if they showed retraction or drifted out of focus. Effective diffusion constants were determined by fitting the fluorescence decay data from the photoactivation experiments using a one-dimensional diffusion model function for FDAP according to^34^. All data sets were tested for normality using the D’Agostino-Pearson and Shapiro-Wilk tests. Statistical outliers were identified using the ROUT method. Equality was assessed using a F-test to compare variances. To compare different concentrations, statistically significant differences from control were determined by a one-way ANOVA with Dunnett‘s post hoc test. All statistical analyses were performed using the GraphPad Prism program.

#### Biochemistry

Tubulin polymerization assays were performed in PEDTA (10 mM NaP_i_, 1 mM EDTA, 0.1 mM GTP pH 7.0) or GAB (10 mM NaP_i_, 3.4 M glycerol, 1 mM EGTA, 0.1 mM GTP pH 6.7) buffers. Assembly time courses of 25 μM tubulin at 37 °C were recorded in an Appliskan (ThermoFisher) by measuring absorbance at 355 nm in the presence of 27.5 μM of compounds **1a, 1b**, PTX, FTX2 or DMSO (as control reaction).

The equilibrium binding constant of **1a** and **1b** to the taxane site of assembled MTs were determined by the displacement of FTX2 from the taxane site in crosslinked MTs^64^. The displacement isotherm of each ligand was measured at least three times in two different plates with a fluorescence polarization microplate reader, as described^65^. A mixture of 50 nM taxane binding sites and 50 nM FTX2 in GAB buffer was sampled in a 96-well plate. Increasing concentrations (12.5 nM – 25 μM) of the competitor ligands (**1a** or **1b**) were added to the wells. After mixing the samples for 10 min by shaking at 250 rpm, they were incubated 20 min at the desired temperatures (26 °C, 35 °C or 42 °C) and their anisotropies were measured using a Spark Tecan plater reader in fluorescence polarization at emission of 535 nm and excitation of 485 nm. A sample containing 50 nM taxane binding sites was employed as blank, and a 50 nM solution of FTX2 in GAB, 0.1 mM GTP was employed as anisotropy standard. G factor was calculated as a mean over several cells in different conditions. The molar saturation fraction, *X_b_*, of the probe for each competitor concentration is calculated from the polarization values measured, *P*. Measurements of the polarization of FTX2 completely displaced (*P_0_*) and, in absence of the competitor ligand (*P_sat_*), are required for data processing. The binding of FTX2 in the presence of the competitor ligand (*X_b_* = (*P*-*P_0_*/*P_sat_*-*P_0_*)·*X_sat_*) was calculated and best fitted to the equilibrium binding constant of the competitor assuming unitary stoichiometry and the binding constant of the reference ligand (FTX2) with a PC program (Equigra v5)^64^.

Once the equilibrium binding constants are measured at different temperatures, the standard free energy of binding at a given temperature is calculated from the binding constant as ΔG_0app_ = −RTLnK_a_, the standard enthalpy of the binding reaction is calculated from the representation of LnK_a_ versus 1/T (Van’t Hoff representation), and the slope of this representation multiplied by −R is equal to −ΔH_0app_. The standard entropy of binding to ΔS_0app_ is calculated from the slope of the linear regression of ΔG_0app_ versus T.

#### Computational model building

*Ab initio* geometry optimization and derivation of atom-centered RESP charges^66^ for FTX2 and **1b** were accomplished using the density functional tight-binding method, a 6– 31G(d,p) basis set, and the IEF-SCRF continuum solvent model^67^ for water, as implemented in program Gaussian 09 (Revision D.01)^68^. The conformational preferences of FTX2 and **1b** in aqueous solution were studied upon immersion of each ligand in a cubic box of TIP3P water molecules and simulation at 300 K using molecular dynamics (MD), as performed earlier for PTX and other taxanes^26^. Bonded and nonbonded parameters for the ligands studied were obtained from the general AMBER force field *gaff2*^69^.

Our starting structure of a straight PF segment consisting of three concatenated α:β dimers (α1:β1-α2:β2-α3:β3) has already been reported in detail^12^. Missing residues 39–48 in the four α subunits were grafted from AlphaFold model AF-P81947-F1-v4 for completeness. For consistency with the Protein Data Bank, residue numbering and secondary structure assignment herein follow the α-tubulin-based definitions^70^. The GTP and GDP molecules in the nucleotide-binding sites of α-and β-tubulin, respectively, were kept, together with their coordinated Mg^2+^ ions and hydrating water molecules. The taxanes studied were best-fit superimposed onto the structure of bound PTX, as reported earlier for similar molecules^12^, to yield the corresponding complexes, which were simulated in explicit solvent at 300 K using the AMBER 18 implementation of the *pmemd.cuda_SPFP* engine and the *ff14SB* force field^71^ for protein atoms and TIP3P water molecules, as previously described.

For the ligand-bound macromolecular assemblies representing three parallel short PFs, each consisting of two concatenated head-to-tail dimers (i.e., (α1:β1-α2:β2-α3:β3)/(α1’:β1’-α2’:β2’-α3’:β3’)/(α1”:β1”-α2”:β2”-α3”:β3”)), use was made of PDB entries 6DPU and 6DPV^72^, which depict a stretch of an expanded or compacted 14-PF MT, respectively, and 3JAK^73^, which corresponds to a symmetrized reconstruction of a compacted 13-PF MT. The molecular graphics program PyMOL (v. 2.6, Schrödinger) was employed for molecular visualization and editing. The Cα trace of PDB entry 3JAK was also used as input to either the PyANM plugin for PyMOL (https://pymolwiki.org/index.php/PyANM) or the elNémo web server^74^ to build an anisotropic elastic network model^75^ for normal mode analysis (NMA) using the default values for cutoff (12 Å), force constant (1.0 kcal mol^-1^·Å^-2^) and maximum displacement (amplitude). The NMA allowed us to characterize the second non-trivial mode as that involved in the [low-frequency/large-amplitude] slow bending motion that drives the cooperative structural fluctuations leading to PF number variation. By saving the coordinates of the model corresponding to the maximum amplitude we obtained a suitable template for an [(α1:β1-α2:β2-α3:β3)/(α1’:β1’-α2’:β2’-α3’:β3’)/(α1”:β1”-α2”:β2”-α3”:β3”)] MT patch with a slightly higher degree of curvature (*i.e*., a smaller radius). Six copies of the refined complex were generated in PyMOL. The (α1:β1-α2:β2-α3:β3) stretch of the first copy was then best-fit superimposed onto the (α1”:β1”-α2”:β2”-α3”:β3”) stretch of the harmonically deformed model. Thereafter, the (α1:β1-α2:β2-α3:β3) stretch of each remaining copy was best-fit superimposed onto the (α1”:β1”-α2”:β2”-α3”:β3”) stretch of the preceding model. This process led quite naturally to formation of a tubular structure consisting of 12 straight PFs (**Fig. S3D**) with one PTX molecule bound in the taxane site of every β subunit. Lastly, the macromolecular ensemble was cleaned up to produce an all-atom model of the MT lattice by (i) removing the redundant subunit copies used for the superposition, (ii) renumbering chains, and (iii) carrying out an energy minimization. For each system studied, the interdimer space was first explored in a minimalist complex (β1:α2) in which all Cα atoms −except those in the more flexible and less well-defined loop regions− were positionally restrained by means of a weak harmonic force constant (2.0 kcal mol^-1^·Å^-2^) to preserve the gross characteristics of each MT subtype. The equilibrated system was then subjected to a simulated annealing procedure, by gradually decreasing the temperature from 300 K to 100 K under the same conditions, and thereafter energy minimized in the absence of any restraints, as reported earlier^12^. The resulting optimized complex representing the interdimer interface was then replicated five times and one copy was superimposed onto the respective pairs in the 2×3 patch of tubulin dimers to produce the corresponding MT piece made up of three PFs, each consisting of two concatenated α:β dimers. Thus, six replicas of each bound ligand and ligand-binding site were sampled simultaneously. For these MD simulations, the GTP:Mg^2+^ bound in α-tubulin was kept whereas GDP:Mg^2+^ was placed in all β-tubulin subunits; in all cases, extra care was taken to keep the distinct hydration shells around the bound cations, as reported in the high-resolution X-ray crystal structures deposited with PDB codes 4I4T and 8BDE.

By copying this 3-PF MT piece and superimposing one lateral PF onto another, one complete turn of a MT was generated (**Fig. S3D**). Since we used three different templates (6DPU, 6DPV and 3JAK), we could build three short optimized MTs with distinct properties that were then subjected to an additional energy refinement using the generalized Born model for the bulk solvent^76^, as implemented in the *pmemd.cuda_SPFP* engine of AMBER. By following this hybrid approach of combining explicit and implicit solvent models within the same simulation, the computational efficiency was increased by more than 100-fold relative to comparable simulations using tens of thousands of water molecules. The atomic models of an expanded PTX-bound 12-PF MT, a compacted FTX2-bound 14-PF MT and a compacted **1b**-bound 13-PF MT have been deposited in the <u>ModelArchive</u> database^77^ with accession codes <u>ma-rkctk</u>, <u>ma-ayvpr</u> and <u>ma-galf8</u>, respectively.

#### TIRF microscopy

Total Internal Reflection Fluorescence microscopy experiments were performed on an inverted widefield microscope Nikon-Ti2 E, equipped with a motorised XY-stage, a perfect focus system, Apo TIRF 60× oil NA 1.49 objective and PRIME BSY (Teledyne Photometrics) camera. MTs were visualized using Interference Reflection microscopy (IRM) and fluorescently labelled proteins were visualized sequentially by switching between microscope filter cubes for GFP and mCherry channels. The microscopes were controlled using NIS Elements software (Nikon). All experiments were conducted at RT. Microscopy chambers were prepared from two sandwiched silanized glass coverslips (22 x 22 mm^2^ and 18 x 18 mm^2^) attached through parafilm strips (heated for 15–20 s at 60°C and gently pressed together) to form ∼2.5 mm wide parallel flow channels. Prior to the chamber assembly, the coverslips were washed in 3 M KOH, cleaned using Zepto plasma cleaner (Diener electronic) and silanized with hexamethyldisilazane. Flow channels were first incubated with 40 µgml^−1^ anti-biotin antibodies (Sigma, B3640) in PBS for 10 min, followed by incubation with 1% Pluronic F127 (Sigma Aldrich, P2443) in PBS for at least 1 hour. F127 was washed with BRB80 (80 mM PIPES, 1 mM EGTA, 1 mM MgCl_2_, pH 6.9).

Porcine brain tubulin was isolated using the high-molarity PIPES procedure^78^ then mixed with Biotin labelled tubulin (Cytoskeleton Inc., T333P) in 1:49 ratio. All MTs were polymerized using 2% biotin-labelled tubulin. PTX post-assembly stabilized MTs (GTP polymerized, then PTX stabilized, stored and imaged in presence of PTX) were polymerized from 4 mgml^−1^ biotinylated tubulin for 30 min at 37 °C in BRB80 supplemented with 4 mM MgCl_2_, 5% DMSO, and 1 mM GTP (Jena Bioscience, NU-1012). Polymerized MTs were centrifuged for 30 min at 18,000g in a Microfuge 18 Centrifuge (Beckman Coulter’s). The pellet was resuspended and kept in BRB80 supplemented with 10 µM PTX. Drug stabilized pre-assembly MTs (polymerized in the presence of drugs and GTP, stored and imaged in presence of drug) were polymerized from 4 mgml^−1^ biotinylated tubulin for 30 min at 37 °C in BRB80 supplemented with 4 mM MgCl_2_, 40 µM drug of interest in 5 % DMSO, and 1 mM GTP. Polymerized MTs were centrifuged for 30 min at 18,000g in a Microfuge 18 Centrifuge (Beckman Coulter’s). The pellet was resuspended and kept in BRB80 supplemented with 10 µM

PTX. GMPCPP stabilized MTs (GMPCPP polymerized) were assembled from 4 mgml^−1^ biotinylated tubulin for 2 h at 37 °C in BRB80 supplemented with 1 mM MgCl_2_ and 1mM GMPCPP (Jena Bioscience, NU-405). Polymerized MTs were centrifuged for 30 min at 18,000g 000g in a Microfuge 18 Centrifuge (Beckman Coulter’s) and pellets were resuspended and kept in BRB80.

Stabilized MTs were flushed into the channel and incubated for 15–30 seconds to allow for surface attachment. Unbound MTs were removed by washing with BRB80 supplemented with 10 μM of drug. Prior the experiment, the solution was exchanged by motility buffer (BRB80 containing 10 mM DTT, 20 mM D-glucose, 0.1% Tween-20, 0.5 mg/mL casein, 1 mM Mg-ATP, 0.02 mg/mL catalase, 0.22 mg/mL glucose oxidase, 10 μM drug). Subsequently, a solution containing either 10.8 nM Kif5b-EGFP or 10/40 nM mCherry-Tau in MB was introduced into the chamber.

To assess kinesin-EGFP motility along MTs, an IRM image (50 ms exposure) was acquired followed by a 1-minute video using 488 nm laser excitation (20% laser intensity, 300 ms exposure) with no delay. Two to three videos were recorded from different fields within the same channel and at least 2 channels were measure for each experimental condition. The same experiment was done in 2 to 3 different days giving an n between 8 and 18.

To monitor Tau-mCherry binding to MTs, 4 min videos were acquired with image acquisition every 7 sec. Two imaging channels were recorded simultaneously, alternating between IRM (50 ms exposure) and 561 nm field (30% intensity, 200 ms exposure). In the first experiment, 10 nM Tau-mCherry was injected immediately after the first frame was acquired. Similarly, a second video of the same area was recorded adding an additional 40 nM Tau-mCherry.

##### Image analysis

All data was collected from at least two independent trials. Unless otherwise stated, all data were analysed manually using Fiji (ImageJ); graphs and statistical analyses were created and performed using GraphPad Prism v.8.0.1. Graphs represent data means, and the error bars represent the standard errors of mean (SEM). Background fluorescence was subtracted from all intensity measurements. For each condition, analyses were performed on 12 randomly selected, in-focus MTs per channel, being identified with IRM. For kinesin motility analysis, MTs were selected in the IRM channel by drawing segmented lines 5 pixels in width. Kymographs were generated using the *KymographBuilder* plugin. Individual motor traces were manually selected as straight lines (1 pixel thick) on the kymographs. From these traces, run length (x-axis), interaction time (y-axis), and velocity (calculated as the ratio of both) were determined using the Multiple Velocity Measurement Tool macro that applies to all ROIs in the ROI Manager of Fiji. Finally, landing rate was assessed by quantifying the number of kinesins bound along each MT and normalizing this value to the total MT length (expressed in μm) and time. To quantify the fraction of the MT surface covered by Tau envelopes, a segmented line with a fixed width of 5 px (0.36 μm) was manually traced along either the envelope-covered regions or the full length of individual MTs. The total length of all envelope segments was then divided by the corresponding MT length to yield the percentage of coverage. This analysis was conducted at multiple time frames: four time points (12 s, 1, 2 and 4 min) in the video recorded under 10 nM Tau-mCherry, and three time points (12 s, 2 and 4 min) for the 40 nM condition. To calculate the fluorescence intensity between envelope-covered and uncovered regions, the intensity of the manually selected fragments was measured. The mean fluorescence intensity of each ROI was calculated by averaging the pixel intensity values weighted by the corresponding area. For each MT, the mean intensity of envelope-covered regions was divided by the mean intensity of uncovered regions, which was also determined by drawing linear ROIs (5 px in width) on non-covered segments of the same MT. To compare the amount of Tau bound per micron of MT across different structural conditions, a segmented line (5 px wide) was drawn along the entire length of each MT. The integrated fluorescence intensity (IntDen) was measured, and this value was normalized to the background and divided by the corresponding MT length (in μm). This analysis was performed at three time points (12 s, 2 and 4 min) in both sequential videos. To calculate the initial velocity of Tau coverage, the length (in μm) of MT segments covered by Tau within the first 12 seconds following the injection of 10 nM Tau-mCherry was measured and normalized to the total length of the corresponding MT.

## ACKNOWLEDGEMENTS

This work was supported by Ministerio de Ciencia e Innovación (PID2021-123399OB-I00 and CNS2023-145079 to M.A.O., PID2022−136765OB-I00 to J.F.D., and PID2022-136307OB-C22/AEI/10.13039/501100011033 to F.G); the European Union’s Horizon 2020 research and innovation program (H2020-MSCAITN-2019-EJD: Marie Skłodowska-Curie Innovative Training Networks (European Joint Doctorate)—Grant Agreement No: 860070 – TubInTrain to F. B, A. S, R. B, J. F. D, D. P); the European Research Council (ERC-2022-SYG grant 101071583 ‘TUBULINCODE’ to Z. L.); institutional support from CAS (RVO: 86652036) and Czech Science Foundation (grant 25-17813S) to M. B. and; R. P. O was supported by an EMBO Short Term Fellowship [number 11234] and MOSBR project MOSBRI-2024-296. We thank staff of beamline BL11-NDC-SWEET (ALBA, CELLS, Cerdanyola del Vallès, Spain) for their support; the Imaging Methods CF at BIOCEV, supported by the MEYS CR (LM2023050 Czech-BioImaging); and the CF Protein Production of CIISB, Instruct-CZ Centre, supported by MEYS CR (LM2023042). We also thank Ganadería Fernando Díaz for calf brains supply.

## CONTRIBUTIONS

Conceptualization, M. A. O. and J. F. D. Investigation, F. B., R. P-O., A. S., O. B., B. A-B., D. L-A., R. H., J. E-G., D. O., and F. G. Formal Analysis, F. B., R. P-O., A. S., O. B., J. G-A., F. G., and M. A. O. Methodology, O. B and J. F. D. Writing – Original Draft, M. A. O. Writing – Review & Editing, M. A. O., D. P., R. B., Z. L., M. B, F. G., F. B. and J. F. D. Funding Acquisition, M. A. O., J. F. D., R. B., D. P., Z. L., F. G., and M. B. Resources, W-S. F., F. G., R. B., D. P., M. B., Z. L. Supervision, M. A. O., D. P., R. B., J. F. D., M. B., Z. L., and V. P.

