## Supplemental_figures for "Dissecting structural and functional determinants of microtubule stabilization through guided chemical modulation"

Supplemetary Figures S1 – S3.

Supplementary Tables S1 and S2.

Supplementary Movies S1 – S17.

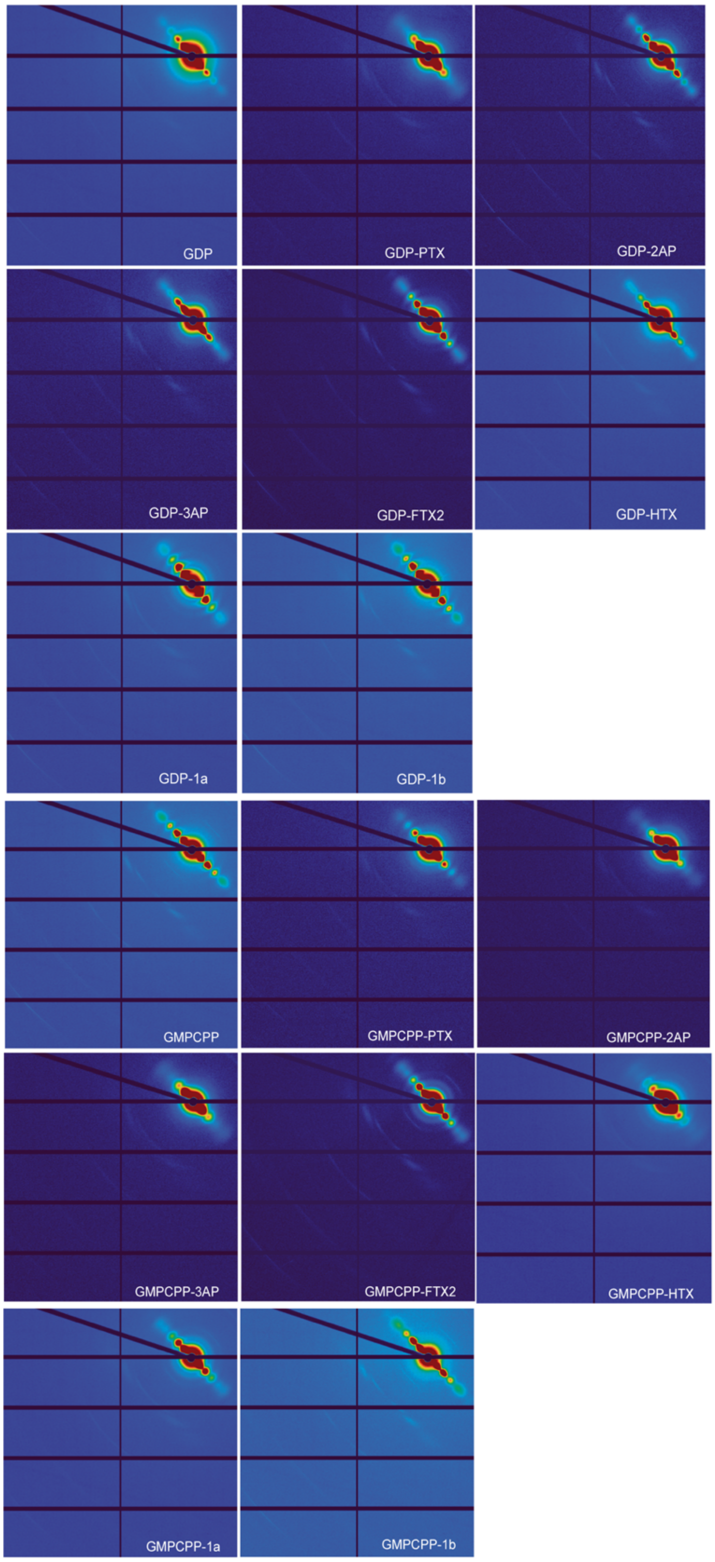

**Figure S1**: **Shear-flow aligned fiber diffraction images** under all conditions analyzed in this study.

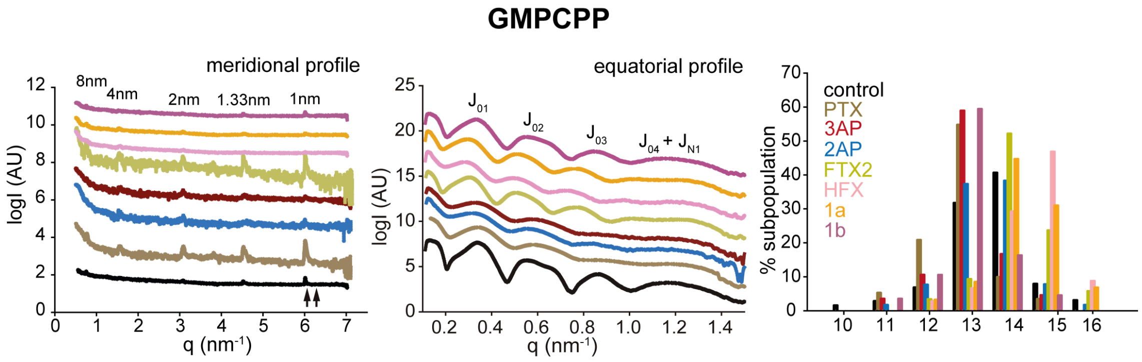


**Figure S2: Fiber diffraction profiles of GMPCPP assembled MTs**. Meridional (left) and equatorial (middle) with and without the different compounds. Profiles are shifted in the y axis for clarity and Bessel functions are labeled. In the left panel, arrows point to the two possible positions of the 1 $nm^{-1}$ layer band (compact vs. expanded lattices). On the right, distribution of MT subpopulations according to the number of PF in MTs assembled with GMPCPP and MTs further stabilized with the compounds used in this study.

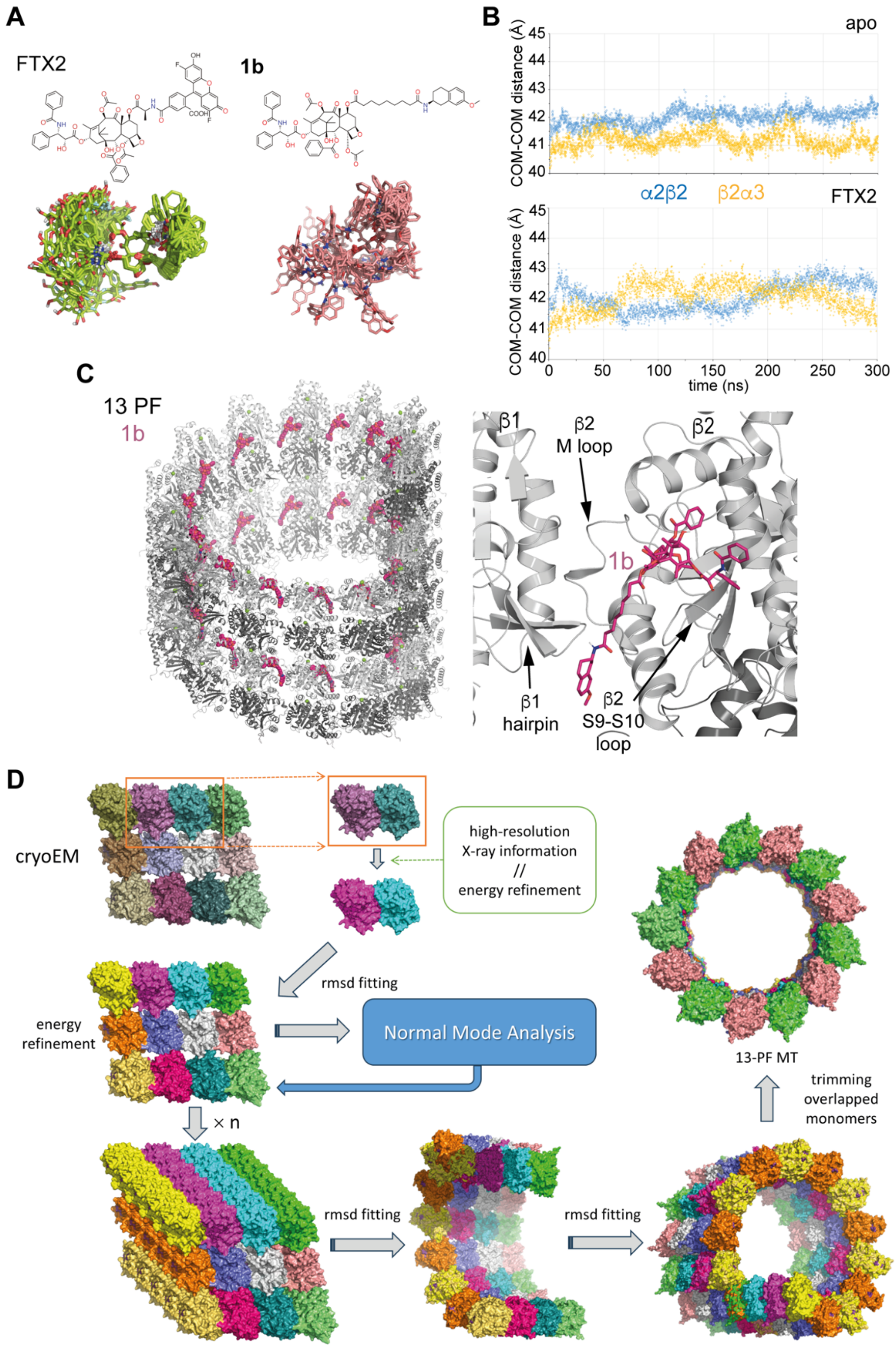


**Figure S3: Computational analysis. A.** Chemical structure (top) and overlay of 3D structural conformers (bottom) of isolated compounds FTX2 and **1b** simulated in a bath of explicit water molecules (waters have been removed for clarity). **B.** Evolution of the distance between α2-β2 (intradimer, blue) and β2- α3 (interdimer, yellow) in the 300-ns MD simulation of the PF both in the absence of ligand at the taxane site (Apo) and in the presence of FTX2. **C.** 3D model of a full turn of a 13-PF MT (cartoon representation) containing a **1b** molecule (purple sticks) bound at the taxane site of every beta subunit (left) and detail of the interprotofilament space showing how the bulky moiety attached to the C7 tail is oriented differently to the FTX2 fluorophore (Fig. 1F), reaching a region below the hairpin involved in lateral contacts of the lateral beta-subunit. **D.** Schematic depicting the MT building procedure. Briefly, an interdimer interface from a MT patch of 3 PFs with 2 tubulin dimers each, as present in the cryoEM structure deposited in the Protein Data Bank) is optimized considering atomic information from high-resolution macromolecular crystals and making use of computational energy refinement in the AMBER force field. The refined interdimer interface is placed back into the cryoEM structure using r.m.s.d fitting and the resulting structure is further optimized using AMBER. Then, this optimized MT patch is reproduced n times and, by r.m.s.d fitting of successive patches, the final MT structure is obtained. The overlapped monomers are removed and the final MT turn is also refined in the AMBER force field.

**Table S1: Normalized motor probes to control experiments**

| | **Cy5-KBP** | | **Cy5-DBP** | |
|---|---|---|---|---|
| | Distance (nm) | Speed ($nms^{-1}$) | Distance (nm) | Speed ($nms^{-1}$) |
| **PTX** | 0.80 ± 0.02 | 0.86 ± 0.02 | 0.81 ± 0.02 | 0.96 ± 0.03 |
| **FTX2** | 0.910 ± 0.003 | 0.85 ± 0.01 | 1.06 ± 0.02 | 1.21 ± 0.02 |
| **1a** | 0.92 ± 0.01 | 0.95 ± 0.01 | 0.92 ± 0.01 | 0.98 ± 0.02 |
| **1b** | 0.85 ± 0.01 | 0.87 ± 0.01 | 0.92 ± 0.01 | 1.07 ± 0.01 |

Data are the mean ± SEM values of two independent experiment with two to three different fields analyzed.

**Table S2: Fiber diffraction data analysis under assembly TIRF conditions**

| | Avg. radius[1] | Avg. PF number[2] | 1 $nm^{-1}$ band[3] | Avg. monomer rise |
|---|---|---|---|---|
| $PTX_{pre}$ | 11.33±0.07 | 12.39±0.12 | 6.02±0.01 | 4.17±0.01 |
| $PTX_{post}$ | 11.52±0.01 | 12.69±0.01 | 6.02±0.01 | 4.17±0.01 |

data are Mean±SEM

[1]nm as calculated from fitting $J_{02}$

[2]calculated from interPF mean distance 5.7 nm

[3]2pi nm$^{-1}$ as calculated from 1 nm layer line

### Movies

**Movie S1:** Normal Mode Analysis of a cryoEM patch of 3 PFs with 2 tubulin dimers each.

**Movie S2**: Movement of Kif5B-EGFP along GMPCPP stabilized MTs.

**Movie S3**: Movement of Kif5B-EGFP along PTX stabilized MTs.

**Movie S4**: Movement of Kif5B-EGFP along **1a** stabilized MTs.

**Movie S5**: Movement of Kif5B-EGFP along **1b** stabilized MTs.

**Movie S6**: Binding of 10 nM Tau-mCherry along GMPCPP stabilized MTs.

**Movie S7**: Binding of 40 nM Tau-mCherry along GMPCPP stabilized MTs.

**Movie S8**: Binding of 10 nM Tau-mCherry along PTXpre stabilized MTs.

**Movie S9**: Binding of 40 nM Tau-mCherry along PTXpre stabilized MTs.

**Movie S10**: Binding of 10 nM Tau-mCherry along PTXpost stabilized MTs.

**Movie S11**: Binding of 40 nM Tau-mCherry along PTXpost stabilized MTs.

**Movie S12**: Binding of 10 nM Tau-mCherry along FTX2 stabilized MTs.

**Movie S13**: Binding of 40 nM Tau-mCherry along FTX2 stabilized MTs.

**Movie S14**: Binding of 10 nM Tau-mCherry along **1a** stabilized MTs.

**Movie S15**: Binding of 40 nM Tau-mCherry along **1a** stabilized MTs.

**Movie S16**: Binding of 10 nM Tau-mCherry along **1b** stabilized MTs.

**Movie S17**: Binding of 40 nM Tau-mCherry along **1b** stabilized MTs.
